# Harnessing the Power of Single-Cell Large Language Models with Parameter Efficient Fine-Tuning using scPEFT

**DOI:** 10.1101/2025.04.21.649754

**Authors:** Fei He, Ruixin Fei, Jordan E. Krull, Xinyu Zhang, Mingyue Gao, Li Su, Yibo Chen, Yang Yu, Jinpu Li, Baichuan Jin, Yuzhou Chang, Anjun Ma, Qin Ma, Dong Xu

**Affiliations:** Department of Electrical Engineering and Computer Science, Bond Life Sciences Center, University of Missouri, Columbia, MO, 65211, USA; Department of Biomedical Informatics, The Ohio State University, Columbus, OH 43210, USA; Pelotonia Institute for Immuno-Oncology, The James Comprehensive Cancer Center, The Ohio State University, Columbus, OH 43210, USA; Institute for Data Science and Informatics, University of Missouri, Columbia, MO, 65211, USA

## Abstract

Single-cell large language models (scLLMs) capture essential biological insights from vast single-cell atlases but struggle in out-of-context applications, where zero-shot predictions can be unreliable. To address this, we introduce a single-cell parameter-efficient fine-tuning (scPEFT) framework that integrates learnable, low-dimensional adapters into scLLMs. By freezing the backbone model and updating only the adapter parameters, scPEFT efficiently adapts to specific tasks using limited custom data. This approach mitigates catastrophic forgetting, reduces parameter tuning by over 96%, and decreases GPU memory usage by more than half, significantly enhancing scLLMs’s accessibility for resource-constrained researchers. Validated across diverse datasets, scPEFT outperformed zero-shot models and traditional fine-tuning in disease-specific, cross-species, and under-characterized cell population tasks. Its attention-mechanism analysis identified COVID-related genes associated with specific cell states and uncovered unique blood cell subpopulations, demonstrating scPEFT’s capacity for condition-specific interpretations. These findings position scPEFT as an efficient solution for improving scLLMs’ utilities in general single-cell analyses.

## Introduction

Single-cell sequencing has revolutionized biology and medicine by providing high-resolution insights into the complex roles and interactions of cell types within their native environments [1]. This technology uncovers critical heterogeneities in tissues, such as those observed in cancer and immune responses, advancing personalized medicine [2]. However, technical challenges such as batch effects and biases, complicate the interpretation of single-cell data [3]. Inspired by the success of foundation models in natural language processing, researchers have developed single-cell large language models (scLLMs), such as scBERT [4], Geneformer [5], scGPT[6], scFoundation [7], SCimilarity [8], GeneCompass [9], and scTab [10]. These models leverage large-scale single-cell atlases to embed the essential biological knowledge intrinsic to single-cell data, helping address these challenges and enabling robust downstream analyses.

The embeddings from scLLMs demonstrate promise in a variety of tasks within familiar cellular contexts [11]. However, scLLMs often struggle in out-of-context scenarios (e.g., unseen diseases, new treatments, uncharacterized cell populations, or cross-species applications), where performance can be unreliable, leading to misinterpretations in zero-shot settings [12-13]. Current efforts to address this typically focus on scaling up both model parameters and pretraining data [14], which come with high computational costs and require extensive data collection, limiting accessibility for many researchers with constrained resources. A more practical solution is to finetune the entire scLLM on out-of-context data. However, finetuning overwrites the original model parameters, potentially erasing valuable pre-learned knowledge, reducing adaptability, and increasing the risk of overfitting task-specific data, which is a phenomenon known as catastrophic forgetting [13]. Additionally, full model updates still require substantial computational resources, which remain a significant barrier.

To address these challenges, we propose scPEFT, a framework that integrates Parameter-Efficient Fine-Tuning (PEFT) techniques into scLLMs to calibrate them for specialized use cases. Unlike traditional fine-tuning, which modifies the entire model, scPEFT employs low-dimensional, learnable, and pluggable adapters to customize scLLMs in a separate, reduced-dimensional subspace. The critical role of these proxy adapters is to estimate a ‘model delta’ (standing for changes in model parameters) for context alignment under the guidance of task-specific objective functions and limited custom data. During the adaptation process, the original scLLM parameters are frozen to preserve pre-learned biological knowledge, while only the smaller adapter parameters are updated. This design reduces the complexity of domain adaptation, enabling higher performance with fewer resources than traditional finetuning strategies and zero-shot queries of scLLMs in out-of-context scenarios. Furthermore, the learned model delta integrates biological context into the attention scores of scLLMs, enabling conditional interpretations that align with pre-learned general gene activities and condition-specific requirements.

We validated the robustness of scPEFT across diverse out-of-context scenarios, including disease-specific datasets, cross-species transfer, and under-characterized cell populations. By mitigating the risk of overfitting to noisy or biased task-specific data, scPEFT demonstrates significant performance gains over traditional fine-tuning approaches and zero-shot models. Moreover, scPEFT significantly reduces the need for parameter tuning and GPU memory. Through attention-mechanism analysis, scPEFT successfully identified COVID-specific cell-state-associated genes and distinguished phenotypic subpopulations in CD34+ enriched and Bone Marrow Mononuclear Cell (BMMC) samples. Additionally, scPEFT further improves scLLM performance in domain-specific tasks such as batch correction and gene perturbation prediction. In summary, scPEFT provides a more efficient and effective framework for unlocking the full potential of scLLMs, facilitating broader applications within the single-cell biology community.

## Results

### Achieving superior performance of scPEFT in cell type identification under disease conditions

The design and technical details of scPEFT are presented in the Methods section and **Fig. 1**. To evaluate its adaptability to disease conditions unseen in the pretrained stage of scLLMs, we focused on cell-type identification using a Non-Small Cell Lung Cancer (NSCLC) dataset comprising diverse T-cell subtypes from the tumor microenvironment. scPEFT was benchmarked with scBERT, Geneformer, and scGPT as backbones, comparing their native and fine-tuned performances. A Uniform Manifold Approximation and Projection (UMAP) [15] visualization (**Fig. 2a**) illustrates the behavior of selected cell representations under native, fine-tuned scGPT, and scPEFT, indicating that the native scGPT struggled with out-of-context scenarios, while its fine-tuned model was prone to catastrophic forgetting. The benchmarking results showed poor performance of native scLLMs in identifying out-of-distribution cells, as their pretraining data were derived from normal conditions (**Fig. 2b**). Fine-tuning improved their performance but was computationally intensive, whereas scPEFT further boosted accuracy under identical conditions and 5-fold cross-validation (39.7%-72.6% accuracy improvements with p-value < 0.001 compared to native models, and 4.3%-9.2% accuracy improvements with p-value < 0.05 compared to finetuned models). Notable performance gaps were observed across the tested scLLMs (**Fig. 2b**), with the four adapter types in scPEFT demonstrating varying levels of efficacy (**Supplementary Fig. 1a**), driven by their gene tokenizer designs and pre-trained knowledge. Nevertheless, scPEFT effectively unlocked the full potential of each model. In contrast, two widely used domain-agnostic tools, SingleR [16] and Seurat [17], appeared difficult to accurately assign cell types, highlighting the challenges posed by inter-patient variability in this dataset.

**Figure 1:**
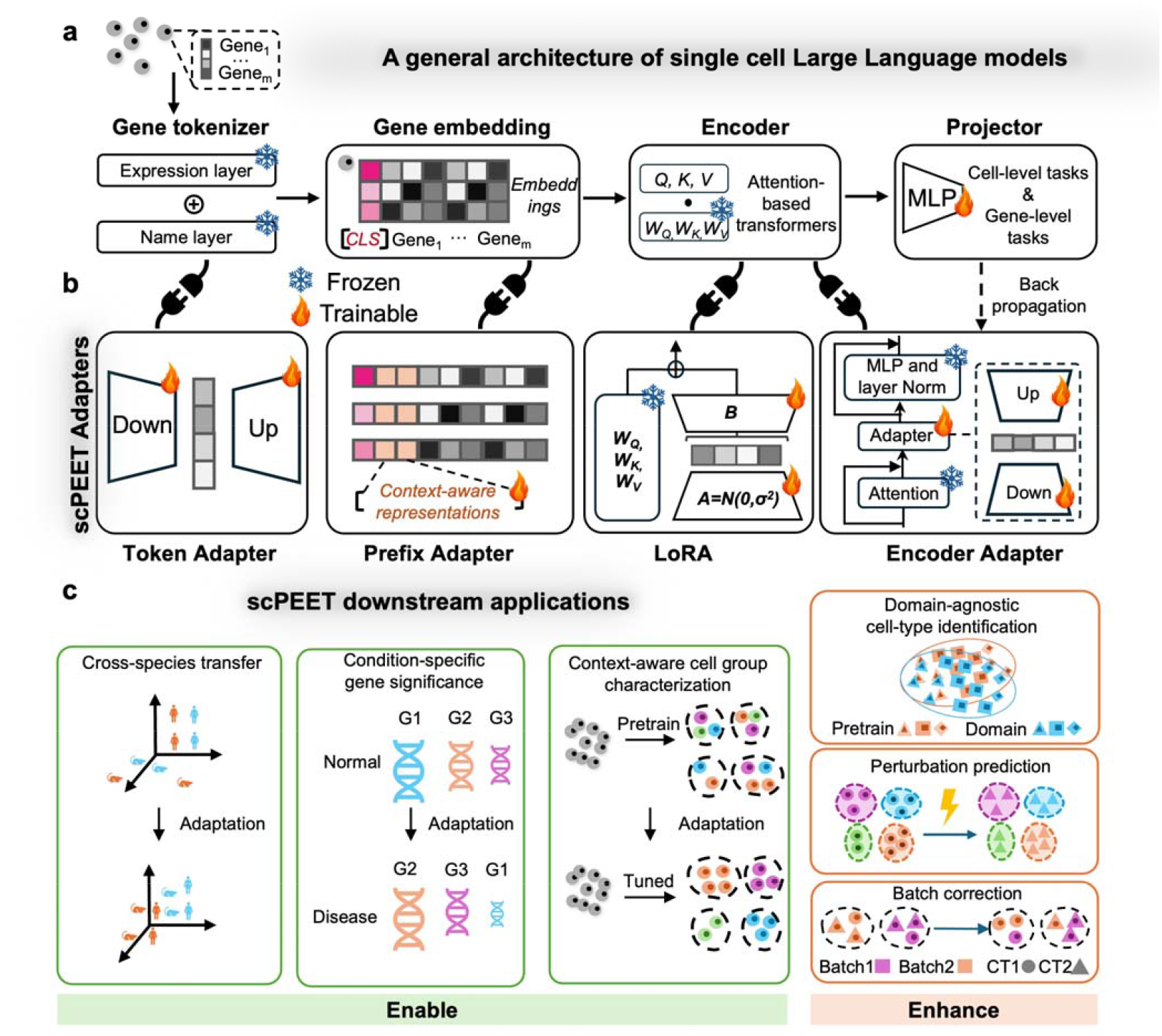
Overview of scPEFT. **a, scLLM Architecture**. A typical scLLM features a gene tokenizer that encodes gene identities and expression profiles into gene embeddings. This is followed by the Encoder, comprising multiple Transformer blocks that aggregate gene expression in cells into gene and cell representations. The final module, a projector, transforms these gene and cell embeddings into task-specific outputs. With the adapters from scPEFT, it can be adapted to various out-of-context applications without updating its original parameters, through task-specific objective functions and back-propagation. **b, Adapters in scPEFT**. Four types of adapters enhance scLLM’s domain adaptability: (i) **Token Adapter**, a compact autoencoder integrated into the gene tokenizer, refines gene token embeddings for specific tasks in reduced dimensional space. (ii) **Prefix Adapter**, which appends tunable tokens to gene tokens to incorporate task-specific information. (iii) **RoLA (Rank-ordered Low-rank Approximation)**, which introduces low-rank matrices A and B into the Transformers to approximate model adjustments for the target domain. (iv) **Encoder Adapter**, another autoencoder attached to a Transformer block, customizes gene contextual embeddings to new biological contexts. These adapters can be used in combination. **c, Downstream applications:** scPEFT tailors scLLMs for a range of downstream applications in specific biological contexts, including domain-agnostic cell-type identification, condition-specific gene significance, context-aware cell group characterization, cross-species transfer, perturbation prediction, and so on.

**Figure 2:**
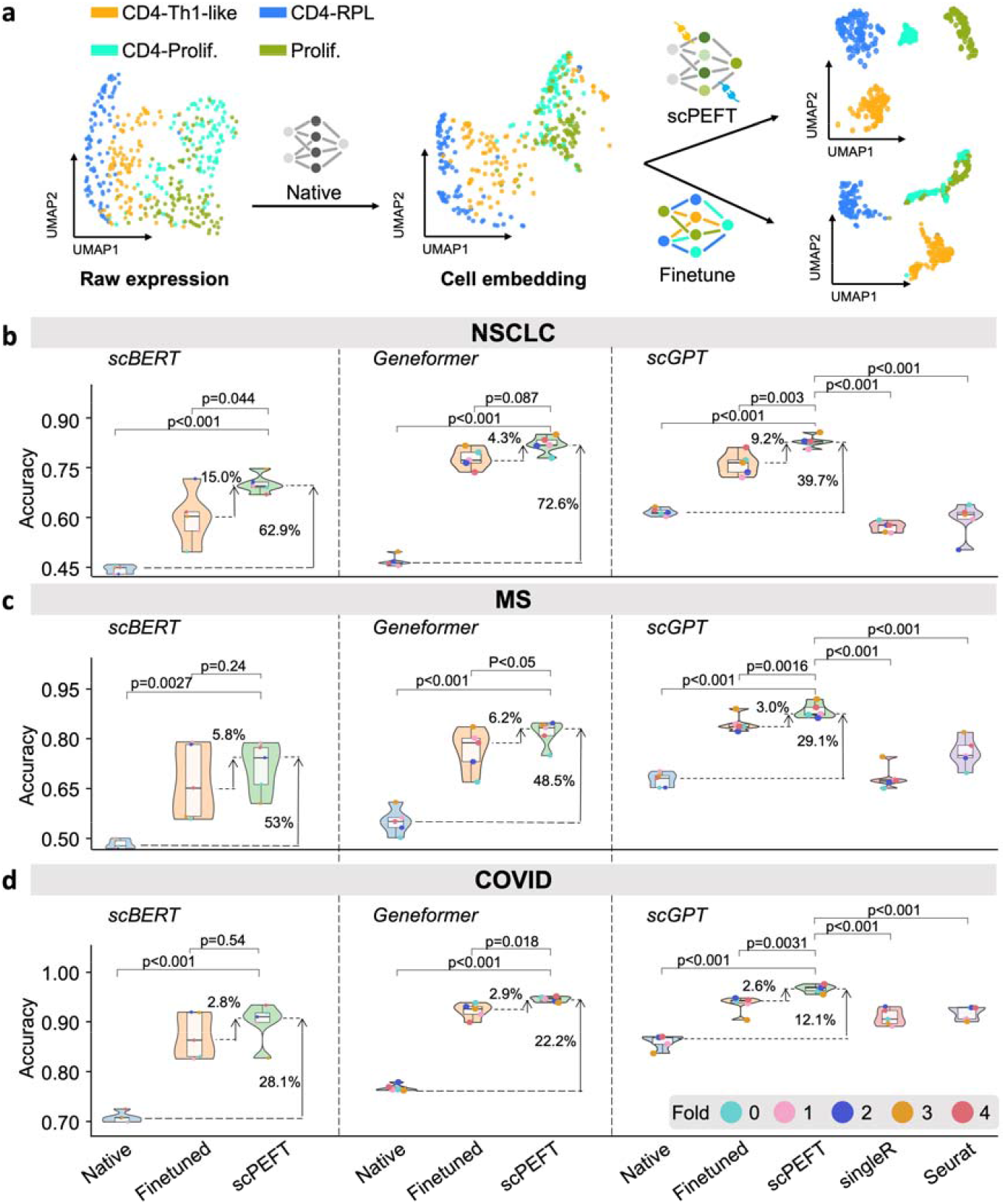
Cell type identification results of scPEFT under disease conditions. **a**, an illustration of how native scGPT, finetuned scGPT, and scPEFT models drive cell embeddings in their feature space. The data points represent a 10% random sample of cells from four annotated cell types in the query partition of the NSCLC dataset. Native scGPT appears to cluster cells according to their identities but misinterprets some CD4-Th1-like cells into CD4-proliferative cells, likely due to expression shifts in the tumor microenvironment compared to its training data of normal cells. Finetuned scGPT achieves better separation of CD4-Th1-like and CD4-RPL cells but performs worse in distinguishing CD4-proliferative cells from proliferation cells than the native scGPT, indicating catastrophic forgetting of pretrained knowledge. In contrast, scPEFT preserves the capability of native scGPT while benefiting from domain adaptation. **b**-**d**, Violin plots benchmarking native, finetuned, and scPEFT models using scBERT, Geneformer, and scGPT as backbones, along with SingleR and Seurat, under five-fold cross-validation on the NSCLC, MS and COVID datasets, respectively. Statistical significance between scPEFT and other models was assessed using a paired Student’s t-test across the five-fold validation results.

Leveraging multiple pretrained checkpoints from scGPT, we replaced its default model with organ-specific variants, including lung-scGPT and pan-cancer-scGPT, as near-context scLLMs for the NSCLC dataset. Interestingly, without adaptation, both lung-scGPT and pan-cancer-scGPT underperformed relative to the default scGPT (**Extended Data Fig. 1**). After applying domain adaptations, scPEFT achieved similar performance across these scGPT variants (39.7%-227.9% accuracy improvements with p-value < 0.001 over native models and 8.5%-10.1% accuracy improvements with p-value < 0.01 compared to finetuned models).

Confusion matrices (**Extended Data Fig. 2**) revealed that fine-tuned scLLMs experienced catastrophic forgetting, failing to recognize certain cell types identifiable by their native models. For example, while the native scGPT model misclassified only 6% of CD4-specific proliferating cells as other proliferating cells, its fine-tuned counterpart showed a 13% misclassification rate. In contrast, scPEFT overcame this performance drop and excelled at identifying rare classes, such as XCL11 cells, which were overlooked by fine-tuned scGPT but successfully identified by scPEFT. These advantages arise from scPEFT’s ability to preserve the original scLLMs’ knowledge, avoiding overwriting during fine-tuning and reducing the risk of overfitting to limited or biased data during domain adaptation. The UMAP visualizations of cell embeddings (**Supplementary Fig. 2**) further exhibit these benefits.

The same comparisons were conducted on the Multiple Sclerosis (MS) and COVID datasets, each holding varying disease conditions. Consistent improvements by scPEFT across the three scLLMs were evident in the performance plots (**Fig. 2cd, Extended Data Fig. 1bc**), while its robustness across diverse cell types and diseases was demonstrated through the confusion matrices (**Extended Data Fig. 3-4**) and the UMAP visualizations (**Supplementary Fig. 3-4**).

### Maximizing efficiency in domain adaptation of scLLMs

We evaluated the computational cost across various adaptation strategies with their default hyperparameter settings (see **Supplementary Table 1**). Comparisons of trainable parameters and GPU memory usage (**Fig. 3a-c**) show that scPEFT utilized only 0.05% to 3.97% of the trainable parameters and less than 50% of the GPU memory compared to fine-tuning scLLMs such as scGPT, Geneformer, and scBERT.

**Figure 3:**
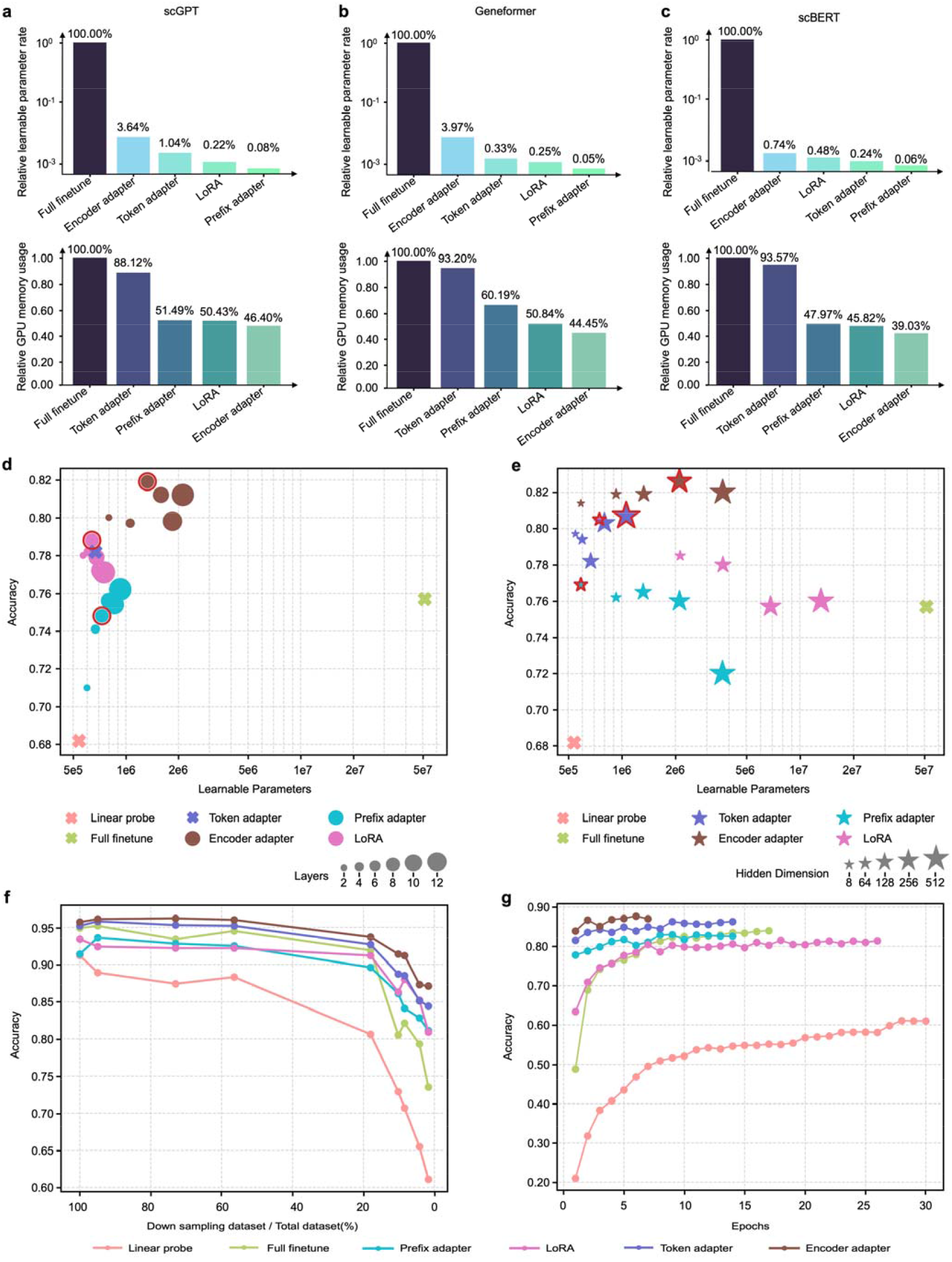
Efficiency analysis of scPEFT and related scLLMs. **a-c**, percentage of learnable parameters and GPU memory usage for fine-tuned scGPT, Geneformer, and scBERT, respectively, relative to scPEFT adapters. Evaluations were conducted using a batch size of 100 cells (the maximum setting for finetuning) on an Nvidia RTX A6000 GPU. GPU memory requirements depend not only on the learnable parameters but also on the gradient propagation path within the model. **d**, validation accuracies plotted against the number of learnable parameters for varying number of Transformer layers with adapters. Larger dots represent configurations with more layers, hence more parameters. Default scPEFT settings are highlighted. Notably, the highest parameter counts may not yield peak performance, suggesting that overparameterization may misalign with the model’s intrinsic dimensionality, thereby reducing generalization, as reflected in lower validation accuracies. **e**, Validation accuracies versus learnable parameters for adapters with different hidden representation dimensions. Larger stars indicate higher intermediate embedding dimensions, requiring more tunable parameters. Default scPEFT settings are highlighted. **f**, validation accuracies of fine-tuned scGPT versus scPEFT adapters using a progressively scaled-down reference dataset. Data points along each curve represent models trained on smaller subsets of the referenced set. **g**, validation accuracies of training checkpoints for fine-tuned scGPT versus scPEFT adapters. The final point on each curve marks the convergence epoch, defined by no improvement in validation loss over five consecutive epochs or reaching the maximum training limit of 50 epochs.

We further assessed scPEFT’s sensitivity to hyperparameter settings. Validation accuracies, calculated using cells held out from a single donor in the reference set, were analyzed in relation to the number of learnable parameters under varying Transformer block configurations on the MS dataset (**Fig. 3d**). Across different hyperparameter settings, most scPEFT strategies consistently outperformed fine-tuning. However, increasing the number of Transformer blocks with adapters did not yield consistent performance improvements. Additionally, we examined configurations with varying intermediate gene and cell embedding dimensions within adapters, observing minimal accuracy variation (<3%) across settings (**Fig. 3e**). Hence, we adopted the default scPEFT settings (**Supplementary Table 1**) for all experiments in this study to ensure generalized results rather than data-dependent optimizations.

We evaluated the impact of annotated data availability across different fine-tuning strategies, recognizing that prior annotations are scarce or costly to obtain. To simulate limited reference data, we randomly reduced the number of donors from reference set while keeping the query set unchanged. The results (**Fig. 3f**) show that as reference data decreased, the validation performance of full fine-tuning dropped significantly, whereas scPEFT remained more robust, indicating that fewer trainable parameters require less data to optimize. We also assessed intermediate checkpoints before early stopping (**Fig. 3g**) and observed that scPEFT strategies consistently achieved higher validation performance, even in the first epoch. In contrast, fine-tuning strategy required more epochs to adjust, showing gradual improvement over time.

### Revealing disease-conditional cell-state-associated genes via scPEFT attention analysis

The attention scores from the scLLMs quantify gene significance in a specific cell group (**Fig. 4a**). scLLMs evaluate individual genes within the context of other genes’ expression, leading to prioritize genes linked to a cell’s phenotype, even if they are lowly expressed or prone to dropout. However, such gene significance interpretation also may stick with their modeling context. We observed the differential attention scores of known signature genes provided by the original study [18] between each T cell sub-type versus other sub-types from native, finetuned scGPT and scPEFT to investigate their ability to spot known cell-type-associated signature genes from the NSCLC dataset (**Fig. 4b**). The attention heatmaps suggested that scPEFT attended these signatures with statistically significant differential attention scores, whereas native and finetuned scGPT tended to miss or misinterpret some of these signature genes.

**Figure 4:**
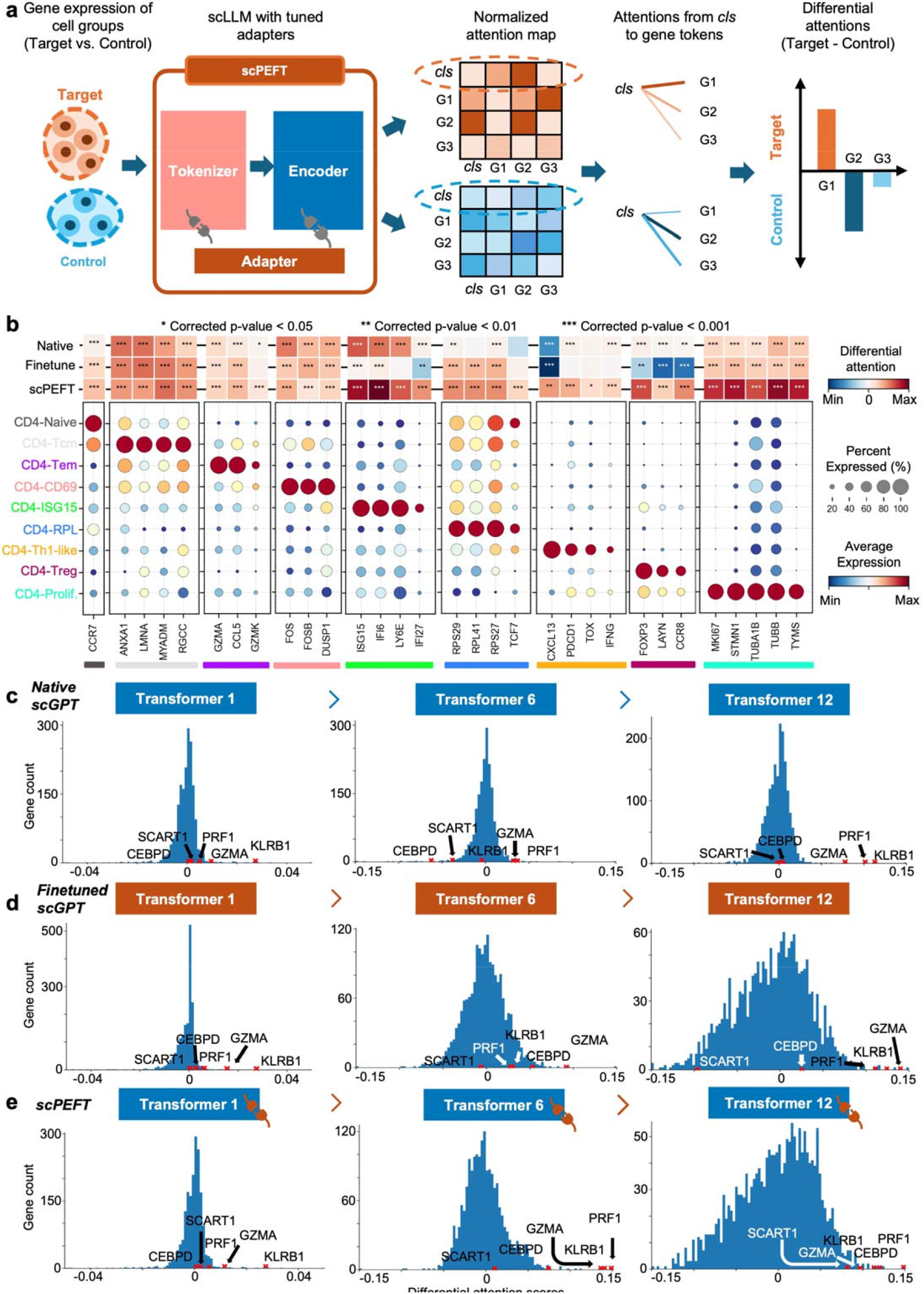
Condition-specific cell-state-associated gene analysis via attention mechanism. **a**, workflow for determining gene contributions to specific cell states under given conditions. Attention scores, describing the cell representation *cls* token’s attention on gene tokens, are extracted and normalized from scLLMs with tuned adapters to assess gene-cell associations. Differential attention scores between control and target cell states are calculated to reveal gene roles in cell-state differentiation under the specified condition. **b**, validation of differential attention values from native and fine-tuned scGPT, as well as scPEFT, in relation to cell-type-associated signature genes on the NSCLC dataset. The cell-type-associated signature genes were sourced from the original study [18]. A heatmap illustrates differential attention scores derived from native, fine-tuned scGPT, and scPEFT models for each signature gene. Differential attention scores were calculated between the target T cell subtype, defined by the corresponding signature genes, and other T cell subtypes as controls. Statistical significance is denoted by stars on the heatmap, based on corrected p-values obtained using the Wilcoxon rank-sum test and Benjamini-Yekutieli false discovery rate control [57]. Dot plots display the expression profiles of each signature gene across all T cell subtypes. Color bars beneath the signature genes indicate their associated target T cell subtypes, corresponding to the subtype labels on the y-axis. **c**-**e**, Histograms of differential attention scores from native, fine-tuned scGPT, and scPEFT models, respectively, in the analysis of COVID-related cell-state-specific genes in Effector Memory CD8+ T cells versus Memory CD8+ T cells. Histograms were generated for the top, middle, and last Transformer layers in these models. Red crosses mark the bins where key effector molecules (*KLRB1, GZMA, PRF1*) and two effector-function-associated genes (*CEBPD* and *SCART1*) were located.

We further compared the attention views of native, finetuned scGPT and scPEFT and differential expression gene (DEG) analysis on different cell states (memory CD8+ vs. naïve CD8+ and effector memory CD8+ vs. memory CD8+) from the COVID dataset. The histograms of differential attention scores (**Fig. 4 cde, Extended Data Fig. 5**) indicate that most genes exhibit minimal differential attention scores in the top Transformer layer, progressively displaying more diverse patterns of differential attention in the middle and last Transformer layers. This progression highlights the capability of stacked Transformers to effectively capture gene-contextual features. Additionally, the greater dispersion of histograms in finetuned scGPT and scPEFT than the native scGPT suggests that these adapted models may provide unique insights into gene-cell associations under disease conditions relative to normal conditions. Conversely, the DEG volcano plots (**Extended Data Fig. 6**), colored by the correlated differential attention scores of these models, show that scPEFT and native scGPT are typically highly correlated with significantly differentially expressed genes identified through DEG analysis. In contrast, the finetuned scGPT appears to overemphasize a broader range of genes across both positive and negative logFC regions. These observations suggest that scPEFT may achieve a better balance between conditional sensitivity and cell-state specificity.

We further examined the genes annotated with the highest attention differences. In the comparison of effector memory CD8+ vs memory CD8+ T cells (**Fig.4cde**), all three models successfully identified three key effector molecules (*KLRB1, GZMA*, and *PRF1*). However, scPEFT uniquely enriched *CEBPD* and *SCART1* with high attention scores, which have been implicated in effector function and tissue-specific homing in T cells [19-21]. These findings suggest that scPEFT may uncover previously uncharacterized regulators involved in the transitions of effector memory T cells. In the comparison of memory CD8+ versus naïve CD8+ T cells from COVID infection (**Extended Data Fig. 5**), *CCL5* and *GZMK* reversed their positions from native scGPT to scPEFT, indicating scPEFT’s recognition of decreased reliance on *CCL5* for memory homeostasis and increased likelihood of an exhausted memory phenotype with increased attention on *GZMK*. Additionally, *CST7* had high attention in only scPEFT, which has been previously associated with transition to effector memory from naïve T cells [22]. These findings reveal that scPEFT provides a valuable alternative approach for identifying disease-conditional cell-state-associated genes.

### Identifying cells from other animals with scPEFT and human data-pretrained scLLMs

Most published scLLMs are pre-trained on human scRNA-seq data. However, building separate scLLMs from scratch for each species is impractical. Given that many cellular mechanisms and functions are conserved across human and other animals [23], we explored adapting scPEFT using orthologous gene for cross-species studies (**Fig. 5a**).

**Figure 5:**
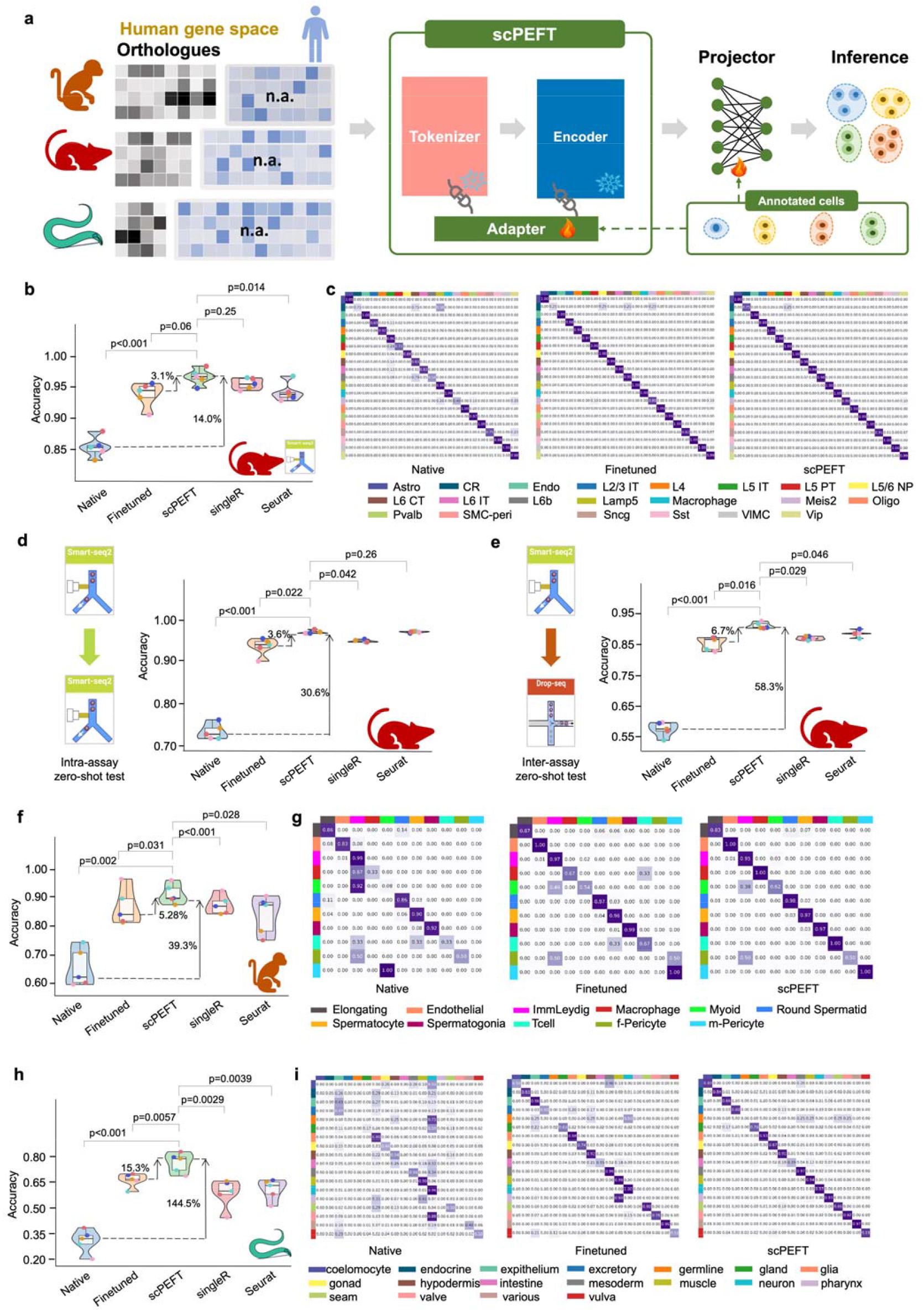
Cross-species transfer results of scPEFT. **a**, Schematic workflow for adapting the human-pretrained scGPT model to data from other species, including mice, macaque, and *C. elegans*. During adaptation, non-orthologous genes are masked, and adapters from scPEFT are trained using a small subset of annotated cells from the target species, enabling cross-species cell type contextualization. **b**, Benchmarking performance of native scGPT, fine-tuned scGPT, and scPEFT across five-fold cross-validation on a mice dataset, alongside comparisons to established cell-type identification tools SingleR and Seurat. Violin plots display performance metrics, with paired Student’s t-tests evaluating the statistical significance of differences between scPEFT and other methods across cross-validation results. **c**, Confusion matrices illustrate the alignment between annotated and predicted cell types for native scGPT, fine-tuned scGPT, and scPEFT on the mice dataset. **d**,**e**, Zero-shot benchmarking violin plots for native, finetune, and scPEFT under five-fold cross-validation on intra-assay and inter-assay interdependent test, respectively, revealing greater stability of scPEFT than finetune strategy in handling assay variance. **f**,**h**, Benchmarking violin plots for native, finetune, and scPEFT under five-fold cross-validation on a macaque and *C. elegans* dataset, respectively. **g**,**i**, Confusion matrices for native, finetune, and scPEFT showing the match between annotated and predicted cell types on a macaque and *C. elegans* dataset, respectively.

Using scGPT as a backbone for scPEFT, we benchmarked the native, fine-tuned scGPT, and scPEFT on a Smart-seq mouse dataset containing finely annotated neuron cell types from the primary visual cortex of healthy adult mice [24]. Results (**Fig. 5b-c**) showed that the native scGPT, using an orthologous gene subset, achieved an accuracy of approximately 75% in identifying mouse cell identities despite not being pre-trained on mouse scRNA-seq data. This suggests the feasibility of adapting human-pretrained scLLMs to identify cell types in other species. Following domain adaptation, scPEFT significantly improved the native model’s performance (14% improvement, p-value < 0.001) and outperformed both the suboptimal fine-tuned model (3.1% improvement, p-value < 0.05) and domain-agnostic tools like SingleR and Seurat.

To examine their applicability, we tested these adapted models in zero-shot settings on two independent datasets with similar cell taxonomies from the same tissue, sequenced separately using Smart-seq and 10x platforms. These tests served as intra-assay and inter-assay evaluations. The fine-tuned model performed consistently in the intra-assay zero-shot test (**Fig. 5d**) but showed a noticeable performance drop in the inter-assay test (**Fig. 5e**), indicating that fine-tuning may be overfitting to custom data, reducing generalizability to unseen datasets with platform-related variations. In contrast, scPEFT maintained robust performance across both zero-shot tests.

We further assessed the adapted models on a macaque dataset containing major germ and somatic cells (**Fig. 5f**). These cell types are rare in the pretraining corpus of most existing scLLMs, posing challenges for the native scGPT in accurately identifying them. After domain adaptation, scPEFT achieved 39.3% improvements in accuracy, indicating its ability to capture shared features between humans and macaques through orthologous genes. In contrast, the fine-tuning strategy demonstrated weaker transferability, as confirmed by confusion matrices (**Fig. 5g**). We also conducted a remote-species test on the *C. elegans* aging atlas. The native scGPT struggled due to the evolutionary distance between humans and *C. elegans* (**Fig. 5hi**). However, scPEFT adapted to achieve accuracies around 80%, representing a 145% improvement over the native model and a 15.3% improvement over the fine-tuning strategy. These results demonstrate that scPEFT regularizes the adaptation process by preserving pretrained knowledge and leveraging human tissue-specific features to form more generalized cell representations for distant species like *C. elegans*.

These experiments demonstrate that scPEFT holds promise for bridging the species divide by capturing a shared embedding subspace of orthologs, facilitating integrated analyses across diverse organisms.

### Uncovering biologically relevant cell populations with scPEFT in an unsupervised manner

Aiming to enable scLLMs to identify new cell states and potentially undiscovered regulators of cell phenotypes and activities from under-studied single-cell data without pre-annotations, we validated scPEFT’s unsupervised adaptation protocol (**Methods**) with scGPT as an example backbone of scPEFT, on a human bone marrow and CD34+ enriched CITE-Seq dataset. CITE-seq protein serves as a natural ground truth for classifying bone marrow cells, which have been extensively characterized using techniques like flow cytometry. Consequently, we annotated these cell identities according to their surface protein expression profile (**Fig. 6ab**) but excluded them from the adaptation process. UMAP visualizations color-coded by each cell’s protein identity (**Fig. 6c**) demonstrated that finetuned scGPT and scPEFT displayed more heterogeneity among the subpopulations compared to that of native scGPT, particularly among granulocytes, B cells, and B cell progenitors. scPEFT demonstrated notable granularity in delineating cellular populations, uniquely identifying nuances such as a subset of mature B cells from BMMC (defined by protein) exhibiting phenotypes closer to progenitor B cells (see the arrow in **Fig.6c**). This may indicate some B cells with remnants of developmental programs not captured by surface proteins or the finetuned scGPT.

**Figure 6:**
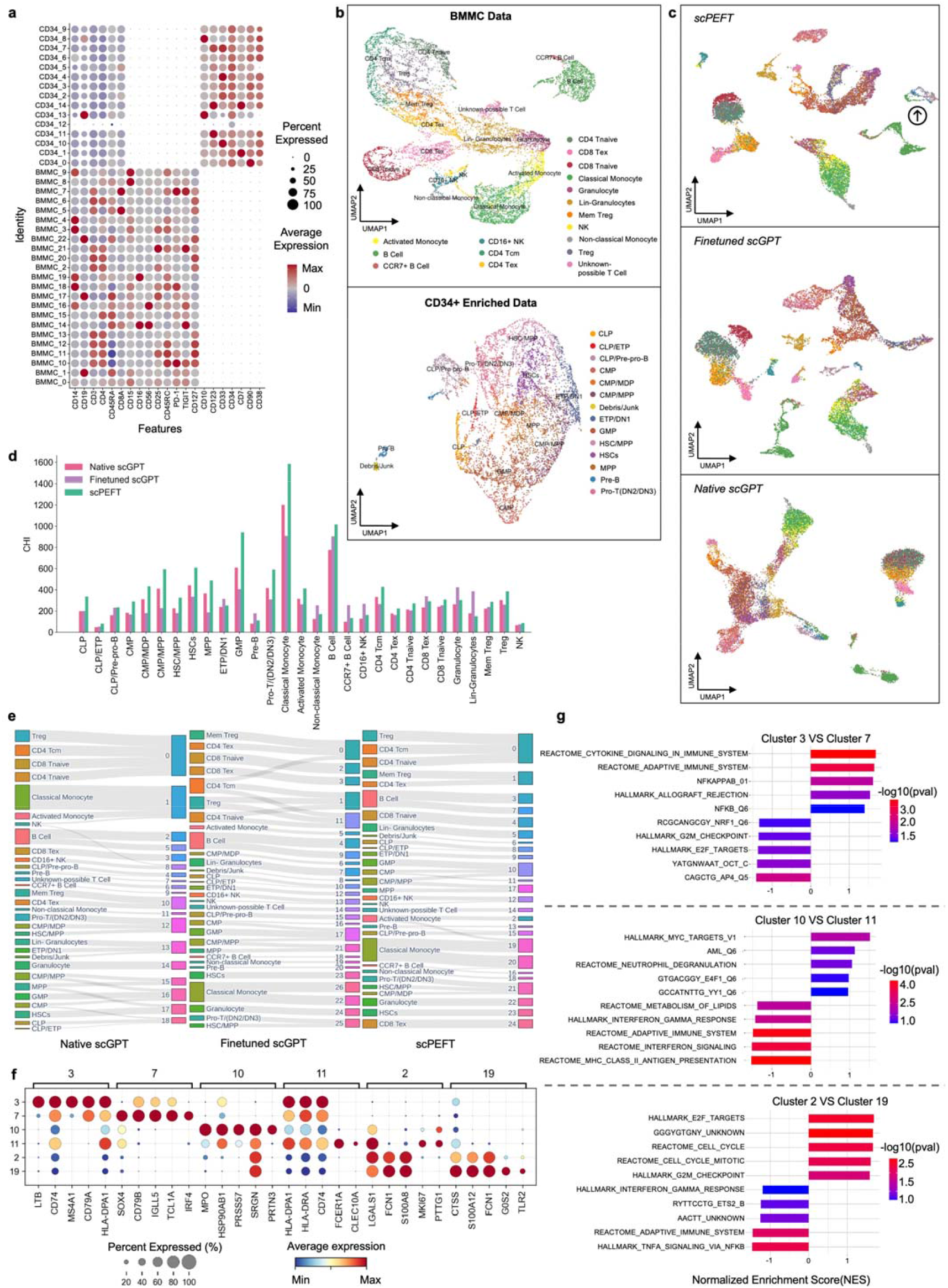
scPEFT identifies developmental cell populations in BMMC and CD34+ enriched CITE-seq data. **a**, Protein expression patterns for annotated cell identities in BMMC and CD34+ enriched samples, depicted by circle sizes representing the fraction of cells expressing a protein and colors indicating average protein expression levels. **b**, UMAP visualizations of protein expression profile clusters from BMMC and CD34+ enriched cells. **c**, UMAP visualizations of cell representations from native, finetuned scGPT, and scPEFT, trained on scRNA-seq data from BMMC and CD34+ enriched cells, excluding cell identity annotations. Arrow points to a BMMC mature B cell subset which clusters closer to pre-proB, and PreB cells, annotated by protein. **d**, Evaluation of cell representations from native, fine-tuned scGPT, and scPEFT models using the Calinski–Harabasz index. This metric assesses the ability of generative embeddings from each model to effectively characterize protein-annotated cell groups. **e**, Sankey diagrams illustrating the assignment of cells from protein-annotated identities (left) to clustering results (right), obtained using the Leiden algorithm at a resolution of 1.5. Clustering was performed on embeddings generated by native, fine-tuned scGPT, and scPEFT models. Finetuned scGPT and scPEFT identified more distinct clusters than native scGPT. Notably, scPEFT was able to avoid some perplexing splitting/combining that confounds the interpretation of the finetuned model. For instance, the finetuned model nearly equally split the *CD4 T*_*cm*_ population between a group of memory *T*_*reg*_ and exhausted CD4 T cells as well as two groups of *T*_*reg*_ and *CD4 T*_*naive*_, combined, suggesting some confusion among memory programs. **f**, Expression profiles of genes receiving high differential attention scores in subgroups #3 vs. #7, #10 vs. #11, and #2 vs. #19 from panel **e**, with colors representing mean expression within each cluster and dot sizes indicating the fraction of cells expressing a gene. **g**, Gene Set Enrichment Analysis (GSEA) based on attention differential scores for these subgroups, showcasing Normalized Enrichment Scores with colors denoting the significance of enriched phenotypes.

We also applied the Leiden clustering algorithm [25] with a resolution of 1.5 to cell embeddings from these models. The CHI scores (**Fig.6d**) indicated that most of the clusters from scPEFT are more compatible with the dispersion of protein-annotated cell groups. It is noteworthy that in most groups, their CHI scores from finetuned model obviously dropped from their native counterparts, evidencing catastrophic forgetting caused by the finetuning process. From the connections between the protein annotations and scRNA clusters (**Fig.6e**), we observed that the finetuned scGPT and scPEFT found a larger number of distinct clusters, compared to the native scGPT, and identified similarly unique splits with no heterogeneity among the surface proteins, like B cells and classical monocytes. However, scPEFT was able to avoid some perplexing splitting/combining that confounds the interpretation of the finetuned model. For instance, the fine-tuned model nearly equally split the *CD4 T*_*cm*_ population between a group of memory *T*_*reg*_ and exhausted CD4 T cells as well as two groups of *T*_*reg*_ and *CD4 T*_*naive*_, combined, suggesting some confusion among memory programs.

We investigated the functional interpretations of novel sub-groups (Clusters 3 & 7, 2 & 19, and 10 & 11) identified by scPEFT using Gene Set Enrichment Analysis (GSEA) [26]. Genes with differential attention scores between sub-groups (top-ranked genes in **Fig. 6f**) were submitted to GSEA, revealing subgroup-specific biological functions (top 5 in **Fig. 6g**). Cluster 3 was enriched for *NFkB* target genes and HLA molecules, while Cluster 7 showed enrichment for pro-growth and developmental pathways (*E2F* and *AP4* targets, *SOX4, IRF4*), suggesting the identification of a recently developed B cell cluster and an activated or memory B cell. Activated monocytes split between Cluster 2 on its own or Cluster 19 where it combines with classical monocytes. Cluster 19 shows high attention for classical monocyte markers (*CD14, CD36, ITGAX*) and enrichment of inflammatory and antigen-presentation pathways (*TNF* and *NFkB* signaling, *CTSS, TLR2*), indicative of an intermediate monocyte population. Cluster 2 monocytes had reduced *CD14* attention, increased *MKI67* attention, and enrichment of pro-growth gene sets, suggesting a true activated monocyte population. Clusters 10 and 11 represented a split of common myeloid progenitors (CMPs), with Cluster 11 enriched for class II molecules and antigen-presentation signatures, consistent with a distinct differentiation pathway. Cluster 10 combined CMPs, granulocyte-monocyte progenitors, and multi-potent progenitors, indicating a mix of premature and undifferentiated myeloid cells not distinguishable by protein panel alone. These findings demonstrate scPEFT’s capability to further resolve biologically relevant subpopulations.

### Enhancing scLLMs on domain specific downstream tasks

We further assessed the efficacy of scPEFT on batch correction and gene perturbation prediction, that require task-specific adaptation to scLLMs instead of their zero-shot settings. We took scGPT as backbone and reused the benchmarking datasets from its publication [6]. For the batch correction task, multiple adapter strategies from scPEFT preserved cell type distributions across multiple batches more effectively (3.95%-9.62% improvements in AvgBIO score on PBMC 10k dataset, Perirhinal Cortex dataset and COVID-batch datasets), and showed competitive batch mixing quality than finetuned model (**Extended Data Fig. 7-9**) and another domain specific tool scVI [27]. The dedicated tool Scanorama [28] outperformed only on COVID-batch dataset but at a higher computational cost in inference. Nonetheless, scPEFT provided a more time- and memory-efficient solution for the batch correction task. In the evaluation of perturbation (**Extended Data Table 1**), no significant differences in average PCCs and MSEs over all genes post-perturbation were observed among scPEFT, finetuned scGPT, and a state-of-the-art tool GEARS [29] across these datasets. When examining the metrics for the top 20 genes with the most significant changes per perturbation, scPEFT also yielded comparable performance compared to finetuned scGPT and GEARS, except for the Norman dataset containing two-gene perturbation. Considering the lower modeling computational demands in terms of memory and time, scPEFT offers a practical and user-friendly approach for gene perturbation prediction task.

## Discussion

scPEFT incorporates small, learnable low-dimensional adapters into scLLMs. By estimating only task-specific deltas, scPEFT reduces the need for parameter tuning by 96%-99.5% and decreases GPU memory usage by half compared to traditional fine-tuning, thereby making specialized scLLMs more accessible to a broader community. scPEFT preserves the pre-learned knowledge embedded in scLLMs, effectively mitigating catastrophic forgetting. We recommend using scPEFT to get context-aware embeddings of the custom data from scLLMs for a wide range of downstream tasks. scPEFT is versatile, functioning in supervised or unsupervised modes for datasets with or without pre-annotations. Additionally, attention analysis of scPEFT enables condition-specific interpretations, facilitating more precise and context-aware discoveries.

scPEFT promotes the capabilities of scLLMs beyond their original pretraining corpus. Notably, its plug-in adapters enable scLLMs, initially trained on human data, to recognize gene programs in other species by leveraging orthologous gene subsets. However, one-to-one orthologous mapping is not always feasible. Integrating scPEFT with strategies that use proteome references to translate gene panels from other species into the human gene vocabulary [30] could help address this limitation.

Further optimization and extension of scPEFT are possible. In this study, we evaluated scPEFT using default hyperparameters across various applications. Optimizing these hyperparameters for specific scenarios could potentially enhance performance. Moreover, the four types of adapters in scPEFT can be combined to suit particular use cases. The pluggable adapters in scPEFT are highly adaptable to any open-sourced scLLM. We plan to develop adapters for some of the new scLLMs, such as scFoundation [7] and CSimilarity [8], and support their user groups. Additionally, scPEFT is extensible to interact with foundational models from other modalities, such as proteomics [31] and imaging [32], enabling efficient, comprehensive, and multimodal understanding of cellular behaviors and interactions.

## Methods

### Architecture of scLLMs

A typical scLLM designs a tokenizer to vectorize gene identities and expression values, and a multi-head Transformer encoder to aggregate gene expression in cells into gene and cell representations, followed by a task-specific projector for inference (**Fig. 1a**).

### Tokenizer

Current scLLMs conceptualize single-cell expression profiles as a form of biological language. Similar to LLMs, current scLLMs require tokenization of genes, converting them into vector representations for downstream learning. However, the key difference in scLLMs is that their tokenizer needs to encode both the gene name and its expression profile. Each scLLM maintains a gene vocabulary based on its training corpus, assigning a unique integer identifier, *id(g*_*j*_), to each gene *g*_*j*_ within a given input cell *i*. Consequently, the gene token vector 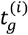 for cell *i* is represented as follows:

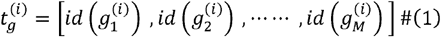

Here, *M* is the total number of genes in the input cell. When input genes do not match the predefined vocabulary, scLLMs omit them from the input. Furthermore, unlike traditional LLMs, scLLMs also incorporate each gene’s expression profile into its token. A prevalent approach utilized by scBERT and scGPT, involves discretizing the normalized expression value *X*_*(i,j)*_ of cell *i* into *m* discrete bins [*b*_1_,*b*_2_,*…,b*_*m*_], yielding the expression profile vector 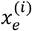:

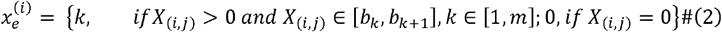

To embed the gene names and gene expression profiles, two embedding layers, denoted as *emb*_*g*_ and *emb*_*e*_ are utilized, respectively, to obtain the final gene representation *h*^*(i)*^ :

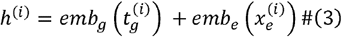

In contrast, Geneformer opts to rank genes according to their relative expression levels across its entire Genecorpus-30M transcriptomic corpus [5], to embed gene expression profile instead. In this case, Equation (2) is defined as

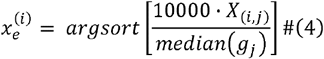

Where *median*(·) retrieves non-zero median expression level of a gene across Genecorpus-30M as a normalization factor, and *argsort* [·] extracts the rank index. The normalized expression values are scaled by multiplying them by 10,000 to enhance precision and are subsequently then adjusted by dividing them by the corpus-wide median normalization factor. This implementation assigns lower ranks to housekeeping genes, which exhibit stable expression, while transcription factors, characterized by greater variability, are ranked higher. This ranking strategy improves the robustness of *x*_*e*_ against technical variants.

### Encoder

The current scLLMs utilize Transformer architecture [33] to model gene expression patterns in cells. This architecture consists of stacking *n* Transformer blocks (*n* = 6 in scBERT [4], *n* = 6 or 12 in Geneformer [5], and *n* = 12 in scGPT [6]), in which each block includes a self-attention layer *Att*(·), two layer normalizations *LayerNorm*(·), and a Multilayer Perceptron (MLP) *MLP* (·). This setup is designed to capture interrelated gene patterns, allowing the computation of all learned gene embeddings 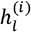 from cell *i* as follows,

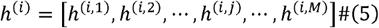

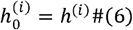

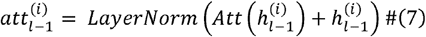

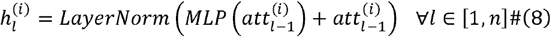

Each cell, represented as a sequence of *M* genes, is then summarized by concatenating or pooling the learned gene-level representations 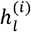 in scBERT and Geneformer, respectively. In certain scLLMs (e.g., scGPT), a special gene-level classification token < *cls* > is placed as *h*^*(i,M+*1)^. This token allows the model to learn an adaptive pooling operation for cell representation through the self-attention mechanism [33] in Transformer blocks.

### Projector

Typically, a Multilayer Perceptron (MLP) serves as a projector, mapping the learned gene embedding 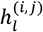 from the last *l*^*th*^ Transformer block into a desired prediction. This gene-level prediction can be a predicted gene expression value or a gene id. For cell-level predictions, the cell embedding *h*^*(i,M+*1)^ is routed through a dedicated projector designed specifically for cell-based tasks. The structure and configuration of these cell projectors are tailored according to the specific cell-level tasks.

### Pretrained settings in scLLMs

In general, scLLMs employ the Masked Language Model (MLM) objective during the pretraining stage to encourage learning of gene contextual features. This objective involves randomly masking selected non-zero gene tokens and predicting the original tokens based on the context provided by the non-masked gene tokens. The learning objective is defined as follows:

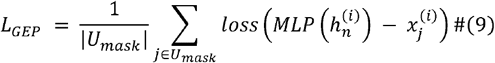

In Equation (9), *U*_*mask*_ represents the set of masked non-zero genes, and 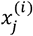 denotes the actual binned or raw expression value for each gene *g*_*j*_ in input cell *i* for scBERT and scGPT, respectively, or the gene identity i Geneformer. The *loss*(·) function measures the difference between the predicted and actual gene information. scBERT and Geneformer utilize cross-entropy [5] loss and scGPT employs Mean Squared Error (MSE) loss [6].

In scGPT, an extra objective models relationships between the special cell token < *cls* > and other gene tokens, defined as

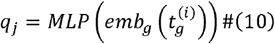

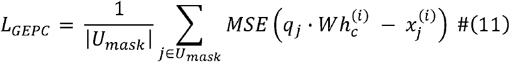

Here, *q*_*j*_ is the gene identity embedding generated by Tokenizer for each gene *g*_*j*_ in input cell *i*. These embedding queries a predicted expression value for gene *g*_*j*_ from cell representation 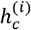, which is the output from the final Transformer block corresponding to the <*cls*> token. The query uses a parameterized inner product *W* with the cell representation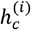, and the mean squared error (MSE) function *MSC* (·) measures the errors between the predicted and actual gene expression values, as shown in Equation (11). During pretraining, an optimizer such as Adam [34] is employed to back-propagate gradients from loss function *L* to update the model parameters. This training process typically utilizes multiple single-cell atlases. For example, scGPT was trained on 33 million cells across various tissues collected from CELLxGENE [35]; scBERT is grounded in the diverse PanglaoDB [36] with over 1.1 million cells; and Geneformer relies on 29.9 million transcriptomes from the Genecorpus-30M [5]. The pretrained scLLMs embed essential biological insights from this extensive corpus and can be used to generate cell or gene representations in a zero-shot setting or serve as a base model for further tuning in diverse downstream applications.

### scPEFT integrates learnable adapters into scLLMs

In scPEFT, we introduce four low-dimensional, learnable, and pluggable adapters into existing scLLMs (**Fig. 1b**). These adapters, with significantly fewer parameters than the scLLMs themselves, are integrated into various components of the models. They are specifically tuned to estimate parameter deltas for their connected scLLM modules, under the supervision of task-specific data and objectives to bridge context gaps and enable diverse downstream tasks in out-of-context scenarios (**Fig. 1c**). During the scPEFT tuning process, the original scLLM parameters remain frozen to preserve the model’s pretrained knowledge.

### Token adapter

Technical variations in single-cell expression profiles across datasets may lead to out-of-context issues. To mitigate this, we developed a tunable adapter, appended to the gene token embedding layer, to calibrate the query gene embedding into pretrained gene embedding space within the tokenizer. This adapter functions as an autoencoder-like module, combining an MLP and a Rectified Linear Unit (RELU) activation to compress *d*-dimensional gene embeddings into a more compact *s*-dimensional format (*s* << *d*). Subsequently, another MLP restores this compressed representation into an adaptive *d*-dimensional gene embedding. As a result, Equation (3) is modified as follows:

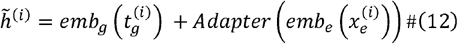

Where *Adapter*(·) denotes the autoencoder-inspired small neural network module. The resulting gene embedding 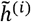 is then fed into the subsequent Transformer-based Encoder. During training, only the adapter parameters are updated, maintaining the native scLLM unchanged. This adapter layer, designed to improve the compatibility of gene tokens across diverse biological contexts, is referred to as ‘Token adapter’.

### Prefix adapter

Extending task tokens to intermediate representations is a popular method for introducing task-specific information in NLP domain. Following the method of Li et al. [37], we insert learnable task tokens to the intermediate embedding, allowing the model to generate task-aware representations. Given the embedding 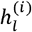 for all genes from cell *i* at the *l*-th Transformer block defined in Equation (5), the extended embedding 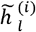 is defined as follows:

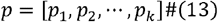

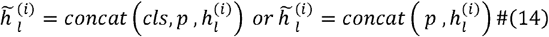

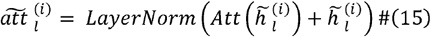

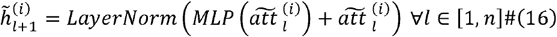

Here, *p* represents the added task token embedding, comprising learnable tensors of the same dimension as the intermediate gene embedding 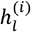 and cell embedding <*cls>* (if applicable for the target scLLM). During training, only the learnable tensors in *p* are updated, while the parameters of native scLLM remain unchanged. This extension, aimed at expanding the semantic capacity of intermediate gene and cell embeddings to incorporate task-specific information, is denoted as the ‘Prefix adapter’.

### LoRA

Low-Rank Adaptation (LoRA) [38] is widely used in NLP foundation models to reduce computational costs during finetuning by introducing low-rank matrices, as illustrated in Figure 1. We employed LoRA technique to efficiently update parameters in self-attention layers of Transformer blocks for downstream tasks. With LoRA, the Quary *Q*_*l*_, Key*K*_*l*_, and Value*V*_*l*_ components in the self-attention layer of *l*^*th*^ Transformer block are computed as follows:

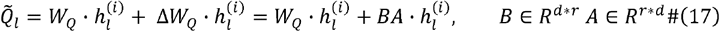

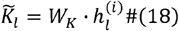

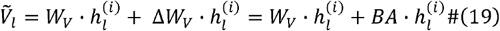

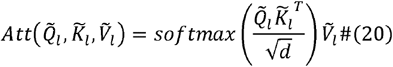

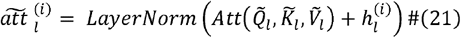

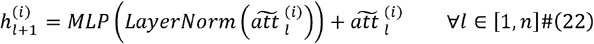

Here, *W*_*Q*_, *W*_*K*_, and *W*_*V*_ represent the pre-train weights for generating Query, Key and Value components, respectively, which remain frozen during finetuning stage to retain the pretrained knowledge. 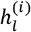 denotes all gene embeddings from cell *i* at the *l*-th Transformer block defined in Equation (5). Updates to the Query and Value components are approximated through two trainable low-rank decomposition matrices,*A and B*, of dimension *r×d* and *r×d*, respectively, where *r* is a predefined low rank (with *r* ≪ *d* and *d* being the dimension of the gene or cell embedding). We only applied LoRA to Query and Value components for simplicity according to [38]. The initialization of *A* and *B* is specified with a random Gaussian and zeros, separately. Hence, *ΔW* = *BA* is zero at the beginning of training on a specific task. After training, task-specific context is embedded implicitly into the low-rank matrices A and B, guiding context-aware gene and cell embeddings 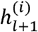.

### Encoder adapter

To further adapt scLLMs for out-of-context data, we integrated an adapter within the Transformer layers in the targeted scLLM, as depicted in Figure 1a. Positioned after the self-attention layers, this adapter aligns learned gene embeddings from context-specific data with pretrained universal patterns, facilitating knowledge transfer from the pretrained model and preventing catastrophic forgetting. In this setup, Equations (7) and (8) are modified as follows:

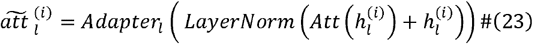

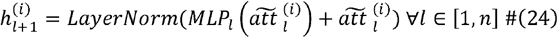

Here, the additional *Adapter*(·) is also the above described autoencoder-like small neural network module. During the training process, only the adapters are updated. We term this component as ‘Encoder adapter’ because it customizes gene and cell relationships within the Encoder module to the specific task, leading to context-aware embeddings that enhance task performance.

### Task-specific learning settings

#### Cell type identification with scLLMs in zero-shot settings

We followed scGPT’s reference mapping workflow to apply scLLMs for cell type identification in zero-shot settings. In this approach, native scLLMs were directly used to embed reference cells and query cells separately. To transfer annotations from the reference set to a query cell, the query cell’s k-nearest reference cells in the embedding space (k = 10 in our study, as suggested in scGPT’s tutorial) were identified. The inferred cell type of the query cell was then determined through a majority vote based on the cell types of these k nearest references.

#### Cell type identification in a supervised manner

To identify cell types, scLLMs typically assign a small multi-layer perceptron (MLP) as a projector and then fine-tune it on a pre-labeled reference set. This projector interprets cell representations (the *cls* tokens or aggregations of all gene embeddings in an scLLM) from the encoder module into categorical cell type predictions. A cross-entropy [6] between predictions and annotated cell types from referenced data was employed as the loss function to supervise training. Accordingly, Equation (9) is adjusted as follows:

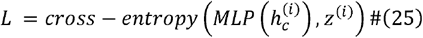

Where 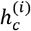 denotes the embedding for the *i*^*th*^ cell, and *z*^*(i)*^ represents its annotated cell type.

We followed the default fine-tuning configurations provided by each scLLM when fine-tuning them for the cell type identification task. During scPEFT’s domain adaptation, we employed the Adam optimizer [34] with an initial learning rate of 10^−^□. Throughout this process, the learning rate gradually decreased to minimize the risk of missing the global optimum. We set a maximum of 100 training epochs and applied an early stopping criterion if there was no improvement in validation loss for 5 consecutive epochs. The tuned model’s performance was evaluated using balanced accuracy as follows under a five-fold cross-validation setting:

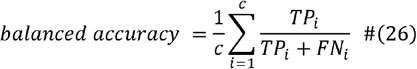

Here, *c* is the number of cell types. *TP*_*i*_ represents true positives for *i*^*th*^ class and *FN*_*i*_ denotes False Negatives for *i*^*th*^ class. This metric treats all cell types equally, regardless of their frequency in the dataset, thus avoiding possible bias in imbalanced datasets. Confusion matrices were also utilized to provide a detailed breakdown of true and false prediction percentages for each class. In the context of identifying *c* cell types, the confusion matrix is a *c*×*c* table where rows represent annotated cell types and columns denote model-predicted cell types. Each cell at *i*^*th*^ row and *j*^*th*^ column reflects the percentage of instances from *i*^*th*^ class predicted as *j*^*th*^ class. Hence, high values along the diagonal indicate accurate predictions, while off-diagonal values represent misclassifications.

#### Attention-based cell-state-specific gene association measure

To quantify each gene’s association with specific cell states, we derive the attention map from the final l^th^ Transformer block as follows:

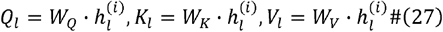

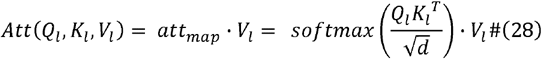

Here 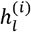 represents the latent representations for gene or cell at the attention layer of *l*^*th*^ Transformer block. *W*_*Q*_, *W*_*K*_ and *W*_*V*_ are the weights from scLLMs using either finetuning or scPEFT. Notably, raw attention scores are computed post-softmax function across multiple attention heads (e.g., 8 heads in scGPT, 10 heads in scBERT, 12 heads in Geneformer, etc). The average over these heads is obtained to offer a comprehensive measurement. In the average attention map, the row indexed by the *cls* token (cell representation) represents the influence of all genes on the cell, as interpreted by the tuned model. This row allows us to observe gene significance within distinct cell groups. Furthermore, to identify genes most strongly associated with a target cell group, we calculated the differential attention score between two closely related cell groups, as illustrated in **Figure 4a**.

#### Orthologous gene mapping for cross-species transfer

To leverage human data pre-trained scLLMs on datasets from other species, we first extracted the overlapping genes between the human gene vocabulary in each scLLM and the custom dataset from other species to define the input feature space. This mapping was implemented using the R package geneSyonym to query the HomoloGene database released by NCBI [39]. Only the matching genes were considered in our cross-species analysis.

#### Cell group discoveries in an unsupervised manner

When prior annotations are unavailable, scLLMs still can be adapted to custom data by reconstructing gene expression levels through self-supervised learning in scLLMs. To this end, some gene expressions are randomly selected to mask. Their predictions are generated from their gene embeddings and *cls* embedding (if applicable) within scLLMs. The pretrained objective functions, such as Equations (11) and (12), are applied in this scenario. In this approach, gene and cell representations simultaneously learn to embed how cell regulatory programs organize themselves and the relationship between this organization and cell activities and phenotypes. The unsupervised tuning of scPEFT was conducted using the same configurations for the optimizer, learning rate, learning scheduler, and early stopping criteria as in the supervised settings. The tuned models produce embeddings for all input cells, which then are clustered using Leiden [40] at a resolution of 1.5 to reveal potential novel cell groups. The dispersion of the annotated cell groups on the generated cell clusters is evaluated using the Calinski-Harabasz Index (CHI) score [41].

#### Batch correction

We employed multiple learning objectives for batch correction as defined by scGPT [6]. Specifically, two pretraining objectives, Equations (11) and (12) are applied to guarantee the context-aware nature of generative gene and cell representations. An additional embedding layer *emb*_*b*_ is incorporated to encode batch information into the learned gene and cell representations from the final Transformer block as:

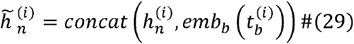

Where 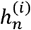 denotes the output gene and cell representations from the final Transformer block, and 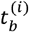 represents the labeled batch identity for cell *i*.

To further enhance batch correction, scGPT introduces an additional objective named Elastic Cell Similarity (ECS), which maximizes the similarity of cell representations sharing the same labeled batch identity. The ECS objective is defined as follows:

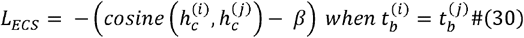

Where 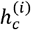 *and* 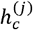 refer to two cell embeddings from a same batch identity. *cosine* similairty is used to measure their dissimilarity, with *β* as a predefined margin.

Another objective, Domain Adaptation via Reverse backpropagation (DAR), addresses the batch artifacts arising from sequencing technology. In this approach, an MLP classifier is assigned to predict the batch identity 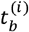 based on each cell representation 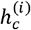.The DAR objective function is as follows

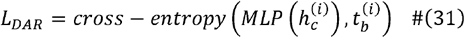

Here, cross-entropy quantifies the error between the predicted and actual batch identity. Eventually, the overall objective for batch correction combines Equations (11), (12), (17), and (18) as shown below.

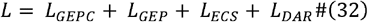

The configurations of the optimizer, learning rate, learning scheduler, and early stopping criteria in this task were set the same as in the supervised settings. For evaluating batch correction performance, we considered two key aspects: biological conservation and batch effect correction, inspired by scIB [42]. Biological conservation is assessed through metrics that evaluate cell-type cluster preservation in the new embedding space, specifically Normalized Mutual Information (NMI) [6], Adjusted Rand Index (ARI) [6], and Cell-type Average Silhouette Width (cell-type ASW) [6]. Batch effect correction is evaluated by examining batch mixing using batch Average Silhouette Width (batch ASW) [6] and Graph Connectivity (GC) [6] as follows:

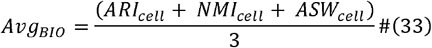

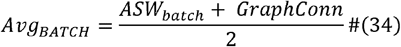

Where Avg_BIO_ donates the average biological conservation score, and Avg_BATCH_ is the average batch effect correction metric.

#### Perturbation prediction

In this task, we used scGPT as the backbone in scPEFT due to its openly available codebase. For perturbation prediction, scGPT considers all genes, regardless of zero or non-zero expression values. To predict absolute perturbed expression levels, we used log1p-normalized expression values as both the input and target, diverging from scGPT’s original design, which used binned values. A binary condition token indicating whether a gene is perturbed was introduced to the tokenizer, modifying Equation (3) as follows:

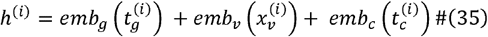

Where *emb*_*c*_ denotes the condition embedding layer, encoding the binary perturbation condition token 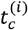 for each gene in cell *i*. This adjustment renders the gene embedding perturbation informed. The objective function for perturbation prediction aims to minimize the difference between the predicted and true post-perturbation gene expression values, defined as:

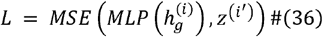

Where 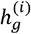 is the gene embeddings of control cell *i* from the final Transformer block, and 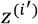 is the perturbated gene expression for a randomly paired perturbated cell *i’* of the same cell type. This approach, aligned with scGPT [6] and GEARS [29], facilitates learning the general perturbated effects across cells. The configurations of the optimizer, learning rate, learning scheduler, and early stopping criteria in this task were set the same as in the supervised settings. Performance was evaluated by comparing predicted and actual post-perturbation expression values using Pearson correlation (PCC) [6] and MSE [6], following the methods of scGPT and GEARS. Additionally, PCC and MSE were calculated for the top 20 most significantly changed genes (denoted as DE_PCC and DE_MSE) to assess the model’s sensitivity to significant expression changes.

### Datasets

#### MS

The MS dataset comprises 21,312 single cells from nine healthy controls and 12 individuals with multiple sclerosis (MS), as annotated in the original study [43]. In alignment with the scGPT data filtering protocol, we excluded B cells, T cells, and oligodendrocyte B cells present only in diseased donors. We selected the top 2,000 highly variable genes per cell for analysis. Cells from 13 random donors were assigned to the reference set, and cells from the remaining 8 donors were assigned to the query set, using a 5-fold cross-validation for performance evaluation.

#### NSCLC

The NSCLC dataset includes 77,030 cells from 36 patients with non-small-cell lung cancer (NSCLC), categorized by treatment response. This dataset emphasizes the role of T cells, annotated by their subtype in the original study [18], within the tumor immune microenvironment. For analysis, cells from 28 patients were randomly assigned to the reference set, and cells from the remaining 8 individuals were assigned to the query set, using 5-fold cross-validation. We selected the top 2,000 highly variable genes per cell to conduct its cell type identification.

#### COVID-19

This dataset consists of 77,168 cells from 16 hospitalized COVID-19 patients with varying symptom severities, including seven asymptomatic, eight moderate, and one severe, as documented in the original study. [44]. We performed a 5-fold cross-validation, randomly selecting cells from 10 patients as the reference set and the remaining six as the query set. The top 2,000 highly variable genes, along with all human transcription factors [45] per cell, were selected for the cell type identification task and gene significance analyses.

#### Mouse

The primary Mouse dataset encompasses 21,874 single cells from two cortical areas of the adult mouse, including the anterolateral motor area (ALM) and the primary visual cortex (VISp), across 127 samples. Four primary and 21 sub-types of cells were annotated in the original study [46]. We randomly partitioned cells from 100 samples to a reference set and cells from 27 samples to a query set under 5-fold cross-validation. The top 2,000 highly variable genes from the orthologous gene set per cell were selected for cross-species validation.

#### Mouse_10x

The Mouse_10x dataset, independently derived from 10x Genomics, encompasses 29,763 cells from ALM and VISp. This dataset was curated from the original study [47] to align cell type contexts with those in the Mouse dataset, thus facilitating a zero-shot query on the models adapted for the Mouse dataset. For consistency and to ensure queries were performed within the same feature space, the same set of 2,000 genes identified in the Mouse dataset was utilized.

#### Mouse_smart-seq

Similarly, the Mouse_smart-seq dataset, derived using Smart-seq v4 technology, includes 15,571 cells from ALM and VISp, sequenced as part of the same study [47] as the Mouse_10x dataset. This facilitates an inter-assay zero-shot query on the models adapted for the Mouse dataset, using the identical set of 2,000 genes to maintain uniformity in the feature space across datasets.

#### Macaque

The Macaque dataset contains 25,481 single cells from five samples, annotated with 11 major germ and somatic cell types by the original study [48]. We randomly reserved cells from one sample as the query set and utilized the remainder as the reference set under 5-fold cross-validation. Similarly, we selected the top 2,000 highly variable genes from the orthologous gene set for model adaptation.

#### C. Elegans

This dataset includes 47,423 scRNA-seq cells from adult *C. elegans*, collected at six different timepoints [49]. Given the species-specific cell type gap, we utilized 19 annotated tissue types as cell identities for evaluation. Cells from one random time point were reserved for query, with the remaining cells used as reference under 5-fold cross-validation. The top 2,000 highly variable genes from orthologous gene set were selected for analysis.

#### BMMC and CD34+

This dataset contains 20,234 BMMCs and CD34+ enriched cells from 4 donors [50]. We used surface protein expression values from CITE-Seq to annotate cell identities based on literature describing populations as ground truth. This resulted in 14 groups representing immune developmental lineages, ranging from HSPCs to Pre-B and Pro-T cells in the CD34+ enriched population, and 17 populations representing largely fully developed immune cells in the BMMCs. This dataset presents a unique challenge for unsupervised analysis, as many of the cells are distinct yet closely linked through developmental lineages. These adjacent cell types differ by the activity of just one or two transcription factors and the expression of as few as one or two surface proteins. The top 2,000 highly variable genes, along with all human transcription factors [45] for each cell, were selected for the cell group discovery and functional analyses.

#### PBMC-10k

This dataset comprises two batches of single-cell RNA sequencing (scRNA-seq) data obtained from peripheral blood mononuclear cells (PBMCs) of a healthy human donor, designed specifically for batch integration evaluation. The first batch contains 7,982 cells, and the second batch includes 4,008 cells, across which nine cell types have been annotated by the original study [51].

#### Perirhinal Cortex

This dataset includes two batches derived from 606 high-quality samples across ten distinct brain regions. The first batch contains 8,465 cells, and the second batch consists of 9,070 cells. Each batch features 10 cell types that have been annotated according to the original study [52].

#### COVID-Batch

The dataset is constructed from 18 distinct batches derived from lung tissue, PBMCs, and bone marrow cells. To ensure a representative cross-section of cell types for robust analysis, it includes only 20,000 selected cells with 39 annotated cell types, as provided by the original study [53].

#### Adamson

This dataset encompasses perturbed gene expression data from the K562 leukemia cell line, generated using Perturb-seq as detailed in the original study [54]. It includes 86 unique single-gene perturbations implemented through CRISPR interference, with each perturbation analyzed in approximately 100 cells. The train set consists of 57 perturbations. The validation set consists of 7 perturbations. The test set consists of 22 perturbations split by GEARS [29].

#### Norman

This dataset features perturbed gene expression data from the K562 leukemia cell line, obtained using Perturb-seq as described in the original study [55]. It is a challenging dataset as it includes 131 two-gene perturbations and 152 one-gene perturbations, with each perturbation replicated in approximately 300 to 700 cells. The train set consists of 137 perturbations. The validation set consists of 30 perturbations. The test set consists of 116 perturbations split by GEARS [29].

#### Replogle_k562

This dataset records genome-wide perturbations in the K562 leukemia cell line, achieved using CRISPR interference as reported in the original study [56]. The dataset is divided into a training set with 278 perturbations and a test set with 103 perturbations by the protocol provided by GEARS [29]. Perturbations without recorded details of the perturbed gene were excluded in our benchmarking, yielding a final count of 68,716 cells across the remaining 412 perturbations.

#### Replogle_rpe1

This dataset records genome-wide perturbations in the RPE1 leukemia cell line, implemented using CRISPR interference as described in the original study [56]. It is divided into a training set containing 439 perturbations and a test set with 163 perturbations, organized according to GEARS [29]. Perturbations lacking records of the perturbed gene were excluded our benchmarking, resulting in a refined dataset of 76,289 cells across the remaining 651 perturbations.

### Benchmarking settings

#### scBERT

We incorporated scBERT [4] as the backbone for scPEFT in our conditional cell type identification and cross-species benchmarking, utilizing its provided cell type identification script with default hyperparameters. Unlike other scLLMs, scBERT standardizes input cell lengths to its gene vocabulary by padding non-existent genes with zero expression values and concatenating all gene embeddings into its projector module. This design results in a significantly larger number of parameters and increased training time compared to other scLLMs. Hence, we implemented the most lightweight Prefix adapter in scPEFT as the default adapter for its benchmarks in this study. Detailed hyperparameter settings are provided in **Supplementary Table 1**.

#### Geneformer

We incorporated Geneformer [5] as the backbone for scPEFT in our conditional cell type identification and cross-species benchmarking, utilizing the provided cell type identification script and the 12-layer version of the pretrained model. As the script includes a hyperparameter optimization process, we reported the optimized results in this study. Given the model’s relative complexity, we adopted the most lightweight Prefix adapter in scPEFT as its default adapter for all benchmarks. Detailed hyperparameter settings are provided in **Supplementary Table 1**.

#### scGPT

We incorporated scGPT [6] as the backbone for scPEFT in our conditional cell type identification, cross-species benchmarking (using its provided cell type identification script), and novel cell group discovery (using its unsupervised pretraining script). In all experiments, we applied the recommended pretrained model trained on the whole human cell atlas, except for the near-context model comparison, where lung- and pan-cancer-specific pretrained models were used. We designated the best-performing Encoder adapter as its default adapter for scPEFT in all experiments. Detailed hyperparameter settings are provided in **Supplementary Table 1**.

#### SingleR

We leveraged SingleR’s reference mapping protocol to benchmark the cell type identification task [16]. The protocol normalizes the input gene expression profiles and then applies Principal Component Analysis (PCA) to reduce the dimensionality of gene expression for both the reference set and the query set. For each query cell, its cell type is assigned based on the most similar reference cells, as determined by their similarities in the reduced-dimensional gene expression space.

#### Seurat

We included Seurat [17] in our benchmarking of cell type identification using its reference mapping protocol. This protocol normalizes expression profiles and represents both reference and query cells through PCA-based low-dimensional reduction. When transferring cell type labels from the reference to the query set, a mutual nearest neighbor strategy is applied. The closest neighbors from the reference set determine the cell type assigned to each query cell. Additionally, we used Seurat’s differential expression analysis script to identify genes contributing to the differentiation of cell types in the COVID-19 dataset.

#### scVI

scVI [27] is a toolbox comprising deep generative models for various omics analyses. It builds a Variational Auto-Encoder (VAE) architecture to model batch effects as latent variables, jointly learning cellular representations to capture biological variability from multiple batches. We benchmarked its batch correction capacity with settings suggested by a benchmarking study scIB [42].

#### Scanorama

Scanorama [28] addresses batch correction by identifying shared cell types across batches through nearest-neighbor searches, utilizing PCA for dimensionality reduction and an approximate nearest-neighbors algorithm based on locality-sensitive hashing and random projection trees. Mutually linked cells create matched batches, which are then corrected and merged into a unified panorama. We also benchmarked its batch correction capability using the settings recommended by scIB [42].

#### GEARS

GEARS [29] is a computational tool designed to predict transcriptional responses to both single and multigene perturbations. By integrating deep learning with a curated knowledge graph, GEARS facilitates the understanding and inference of cellular reactions to genetic modifications for guiding the design of perturbational experiments. We utilized GEARS’s data preprocessing script to prepare data partitions for all benchmarks in the gene perturbation task, adhering to the settings recommended by its official guidelines to evaluate its performance.

## Data availability

All datasets used in this study are publicly available. The NSCLC, BMMC and CD34+,Mouse, Mouse-10x/smart-seq, and macaque datasets can be accessed under number <u>GSE179994</u>, <u>GSE139369</u>, <u>GSE115746</u>, <u>GSE185862</u> and <u>GSE142585</u> of the Gene Expression Omnibus (GEO), respectively. The M.S. dataset is available at https://www.ebi.ac.uk/gxa/sc/experiments/E-HCAD-35. The COVID-19 dataset can be accessed through <u>DOI: 10.6084/m9.figshare.16922467.v1</u>. The *C. Elegans* dataset is available from Calico Research at https://c.elegans.aging.atlas.research.calicolabs.com/. The Adamson, Norman, Replogle_k562 and Replogle_rpe1 datasets can be extracted and preprocessed using GEAR’s *PerData* class *load* function by specifying the *data_name* argument as ‘norman’, ‘adamson’, ‘replogle_k562_essential’, and ‘replogle_rpe1_essential’, respectively. The PBMC 10K dataset can be extracted via scvi’s CLI: *scvi*.*data*.*pbmc_dataset()*. The Prirhinal cortex and COVID-BATCH datasets can both be downloaded from scGPT’s <u>GitHub</u> (https://github.com/bowang-lab/scGPT/tree/main/data).

## Code availability

The source code is freely available on Github (https://github.com/coffee19850519/scPEFT/tree/main).

## Acknowledgements

We thank Mengran Zhao for her technical support. This work was funded by the National Institutes of Health (NIH) R35GM126985. Q.M. is supported in part by NIH R01GM152585, P01CA278732, P01AI177687, U54AG075931, R01DK138504, and the Pelotonia Institute of Immuno-Oncology (PIIO).

## Author contributions

D.X. and Q.M. supervised the study. D.X. and F.H. conceived the study. F.H. and J.E.K. collected datasets and designed case studies. F.H., R.F., X.Z., and M.G. designed the algorithmic framework, analyzed data, visualized experimental results, and developed the software. L.S. and Y.Y. performed experiments and contributed analytical tools. F.H., J.E.K., R.F., and X.Z. wrote the manuscript. D.X. and Q.M. revised and improved the manuscript. Y. Chen, J.L., Y. Chang, and A.M. analyzed and visualized experimental results. All authors approved the final version of the manuscript.

## Supplementary information

**Supplementary Figure 1:** Violin plots of diverse adapters in scPEFT across datasets.

**Supplementary Figure 2:** UMAP visualizations of cell type identification on the NSCLC dataset.

**Supplementary Figure 3:** UMAP visualizations of cell type identification on the MS dataset.

**Supplementary Figure 4:** UMAP visualizations of cell type identification on the COVID dataset.

**Supplementary Table 1:** Default hyper-parameter settings.

**Extended Data Figure 1:**
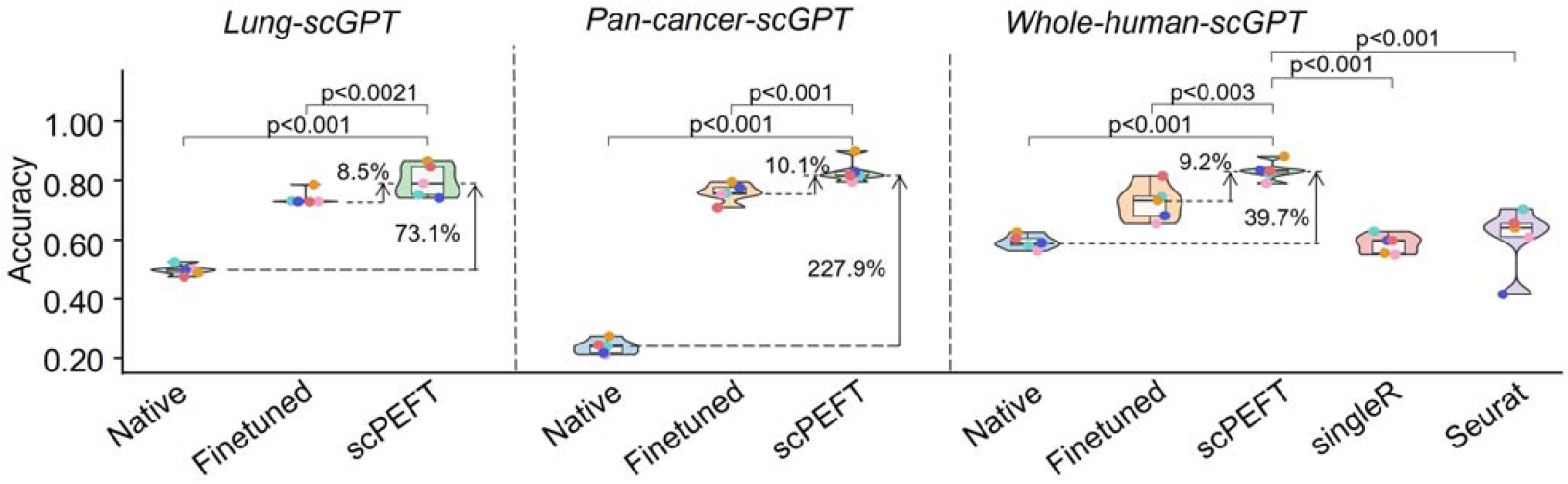
Violin plots of benchmarking results using different scGPT pre-trained weights on the NSCLC dataset. The violin plots benchmark native, finetuned scGPT, and scPEFT using scGPT pre-trained weights from whole human data, pan-cancer data, and lung-specific data.

**Extended Data Figure 2:**
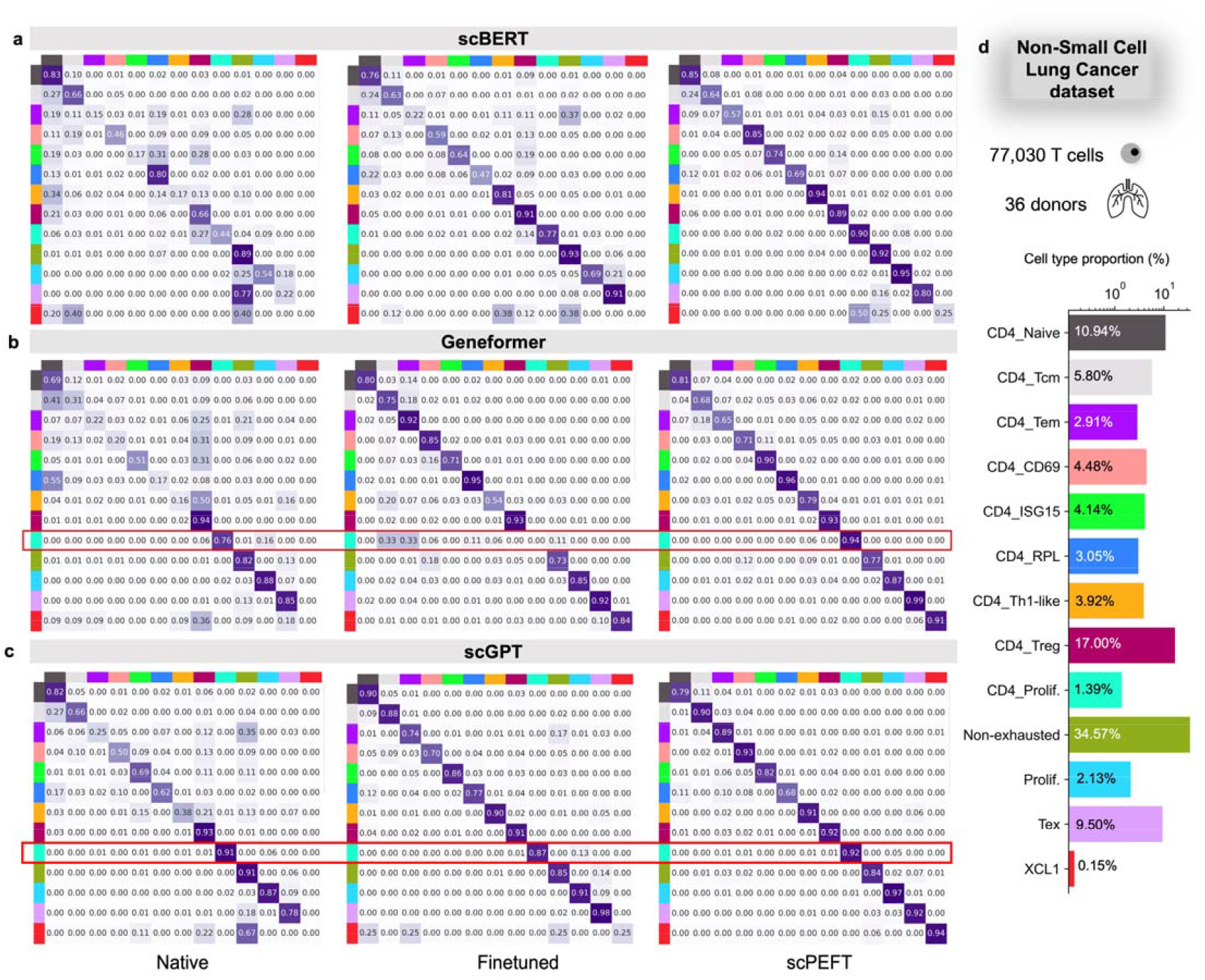
Confusion matrices for benchmarking cell type identification on the NSCLC dataset. **a-c**, Confusion matrices for native, fine-tuned scLLMs, and scPEFT with (a) scBERT, (b) Geneformer, and (x) scGPT as respective backbones. Each cell reflects the percentage of instances from the row-defined cell type that are predicted as the column-defined cell type. High values along the diagonal indicate accurate predictions, while off-diagonal values represent misclassifications. The results exhibit the superior performance of scPEFT, particularly in identifying rare cell types (proportion < 5%). Notably, CD4+ enriched proliferation T cells were accurately identified by native scLLMs but misclassified by finetuned models, demonstrating an instance of catastrophic forgetting. This issue was not observed by scPEFT and was marked by the red boxes. **d**, Bar plot depicting the proportion of each cell type in the dataset.

**Extended Data Figure 3:**
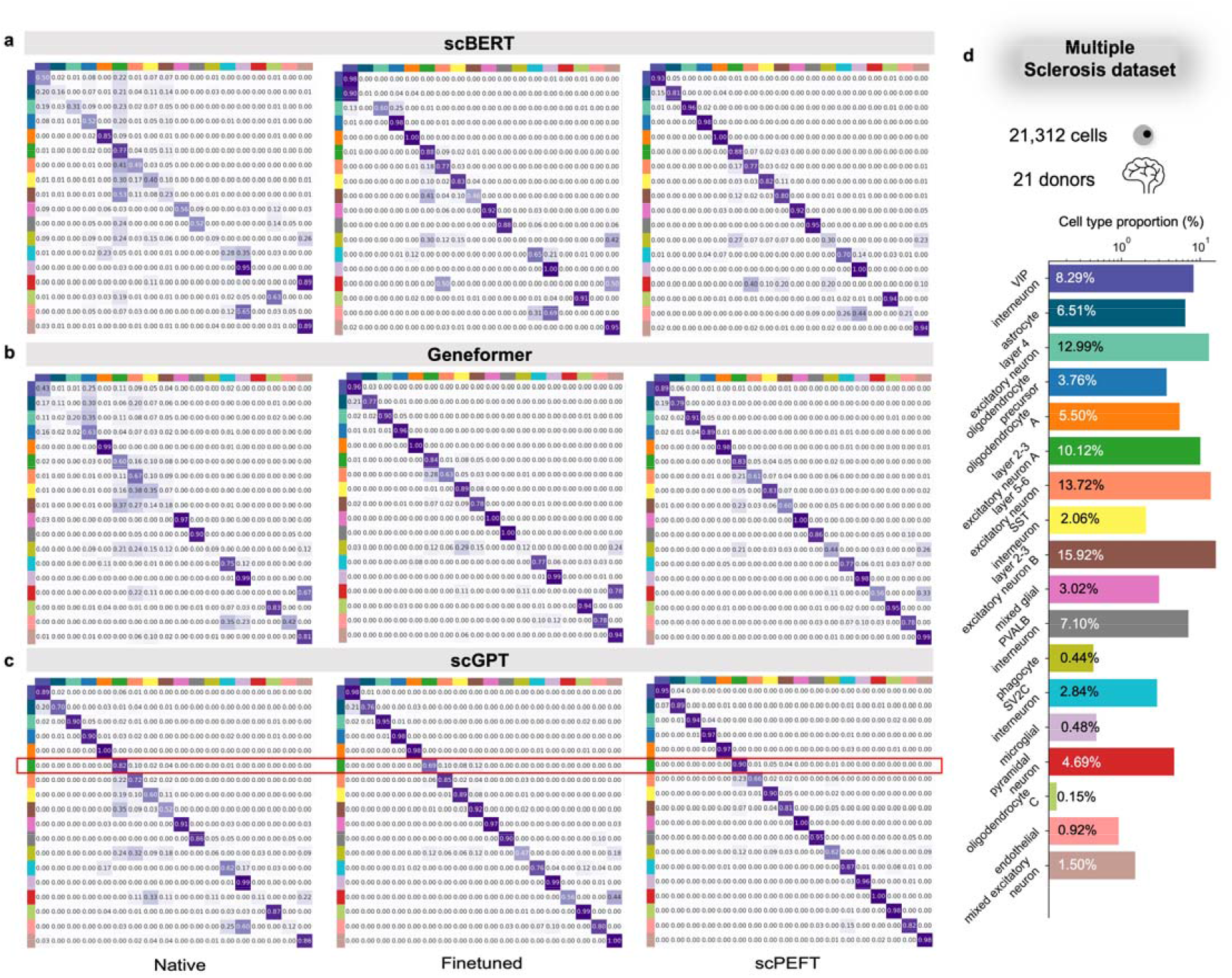
Confusion matrices for benchmarking cell type identification on the MS dataset. **a-c**, Confusion matrices for native, fine-tuned scLLMs, and scPEFT with (a) scBERT, (b) Geneformer, and (c) scGPT as respective backbones. Notably, multiple cell types were accurately identified by native scLLMs but misclassified by their finetuned models, demonstrating clear instances of catastrophic forgetting. This issue was not observed by scPEFT and was marked by the red boxes. **d**, Bar plot depicting the proportion of each cell type in the dataset.

**Extended Data Figure 4:**
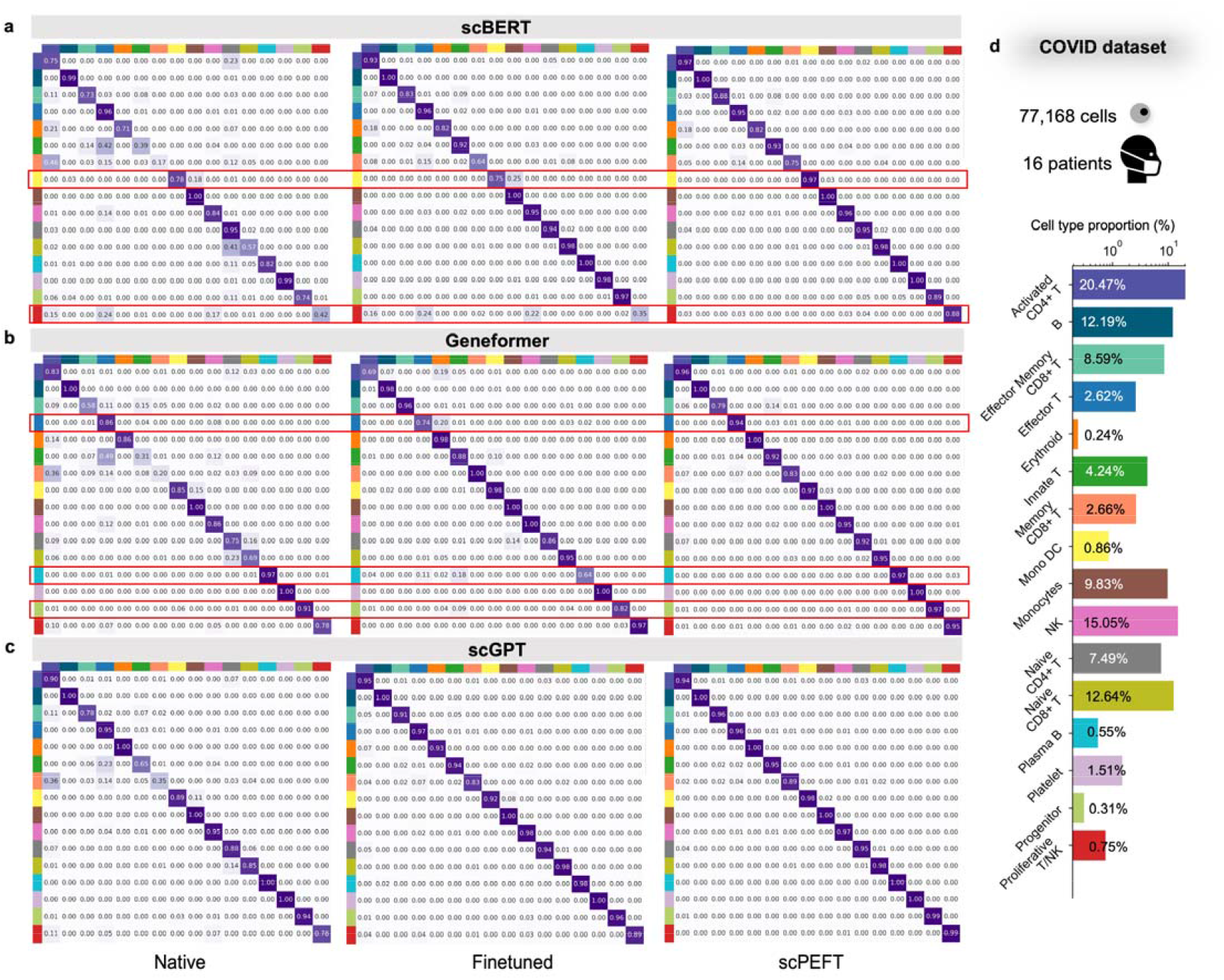
Confusion matrices for benchmarking cell type identification on the COVID dataset. **a-c**, Confusion matrices for native, fine-tuned scLLMs, and scPEFT with (a) scBERT, (b) Geneformer, and (c) scGPT as respective backbones. Notably, the excitatory neuron cells at cortex layer 2-3 were accurately identified by native scGPT but misclassified by its finetuned model, demonstrating an instance of catastrophic forgetting. This issue was not observed by scPEFT and was marked by the red boxes. **d**, Bar plot depicting the proportion of each cell type in the dataset.

**Extended Data Figure 5:**
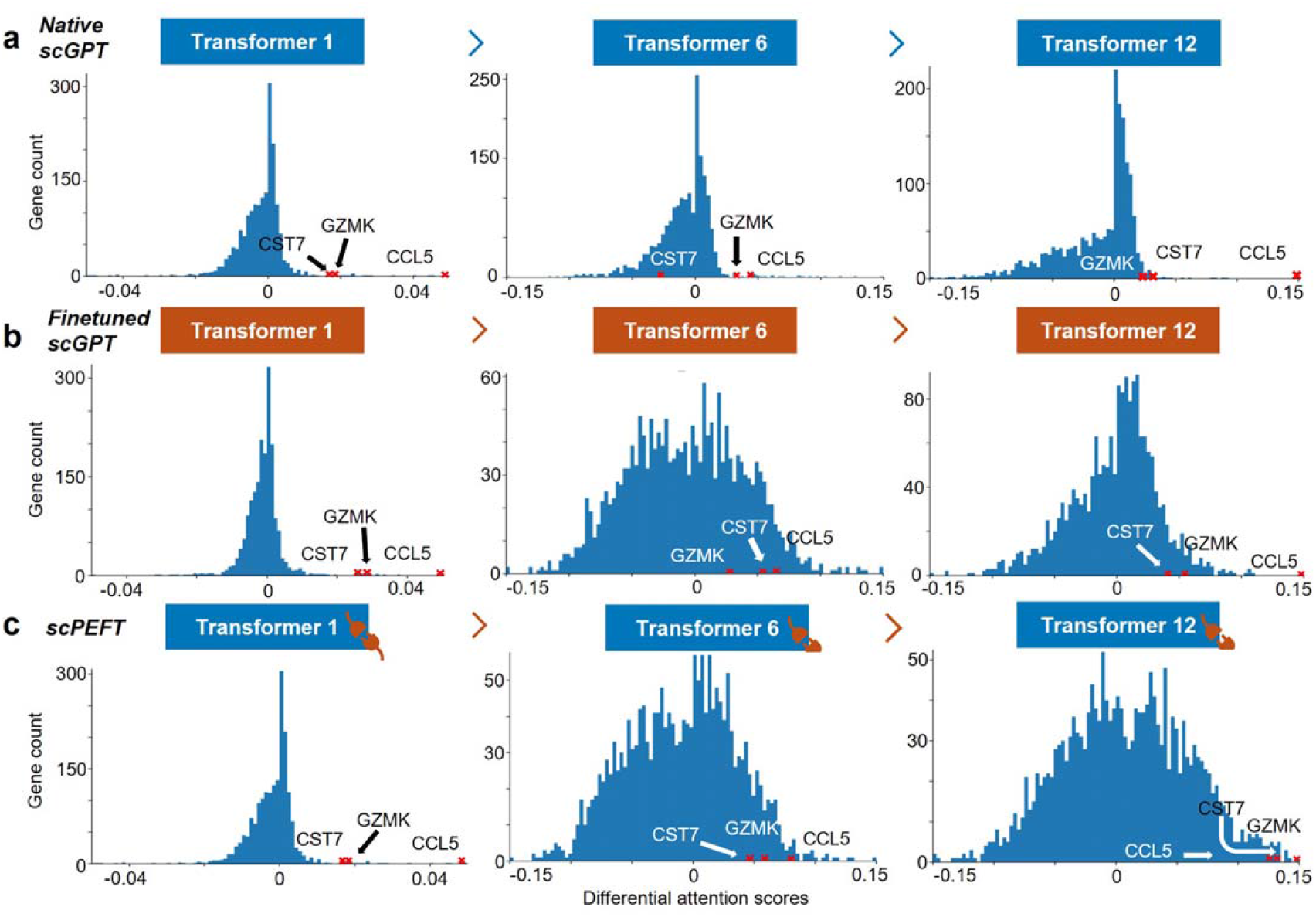
Histograms of differential attention scores for COVID-related cell-state-specific genes. Histograms of differential attention scores were derived from (a) native, (b) fine-tuned scGPT, and (c) scPEFT models, respectively, in analysis of COVID-related cell-state-specific genes in Memory CD8+ T cells versus Naïve Memory CD8+ T cells. Histograms were generated for the top, middle, and last Transformer layers in these models. Red crosses mark the bins where genes of interest (CCL5, GZMK, and CST7) were located.

**Extended Data Figure 6:**
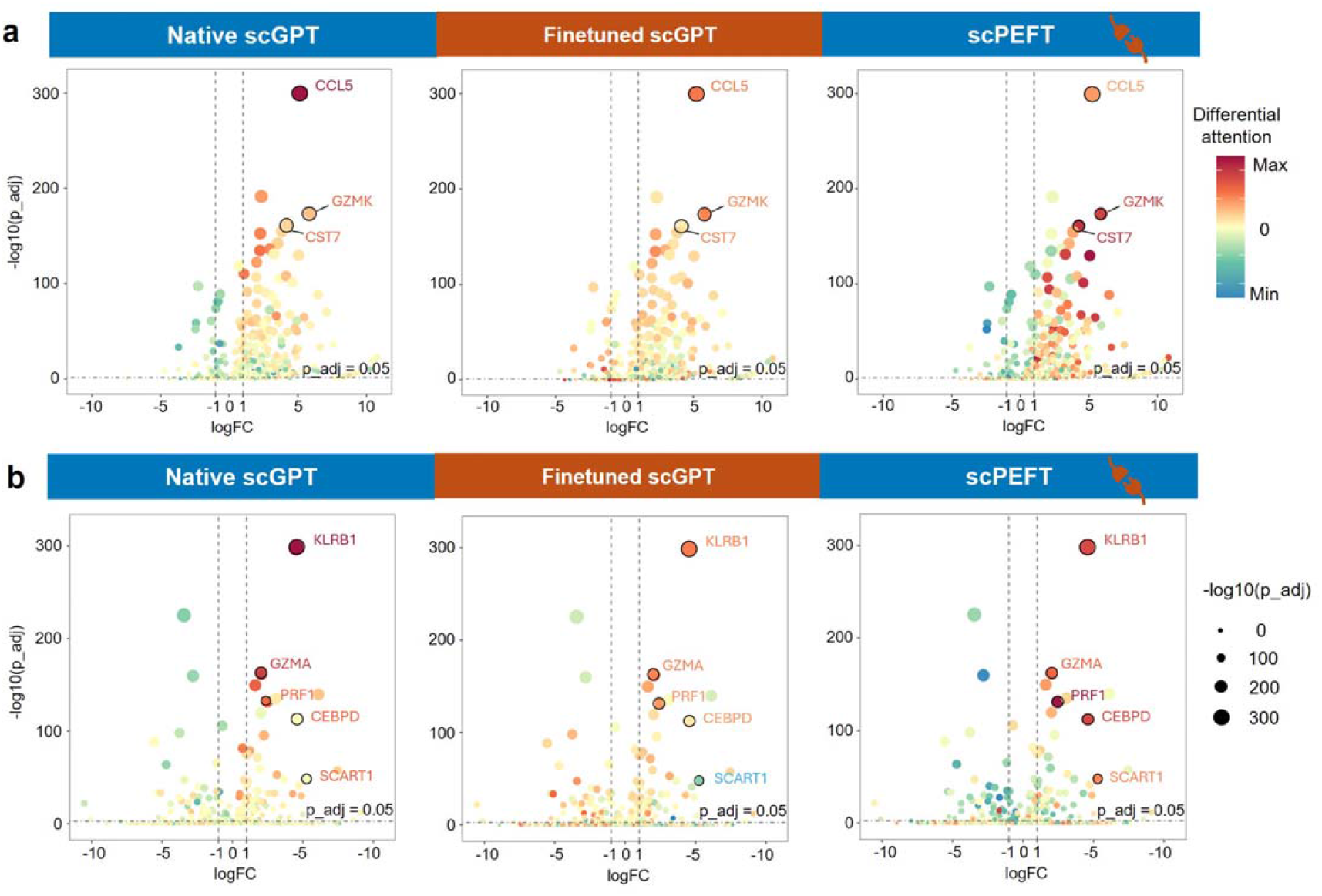
Volcano plots of COVID-related cell-state-specific genes. The COVID-related cell-state-specific genes were analyzed in the comparisons of (a) Memory CD8+ T cells versus Naïve Memory CD8+ T cells, and (b) Effector Memory CD8+ T cells versus Memory CD8+ T cells, respectively. Volcano plots of Differential Gene Expression (DEG) analysis are colored by differential attention scores from native scGPT, finetuned scGPT, and scPEFT trained on the COVID dataset. The top attention genes of interest were highlighted with their names. Dot size reflects the genes’ adjusted p-values from DEG.

**Extended Data Figure 7:**
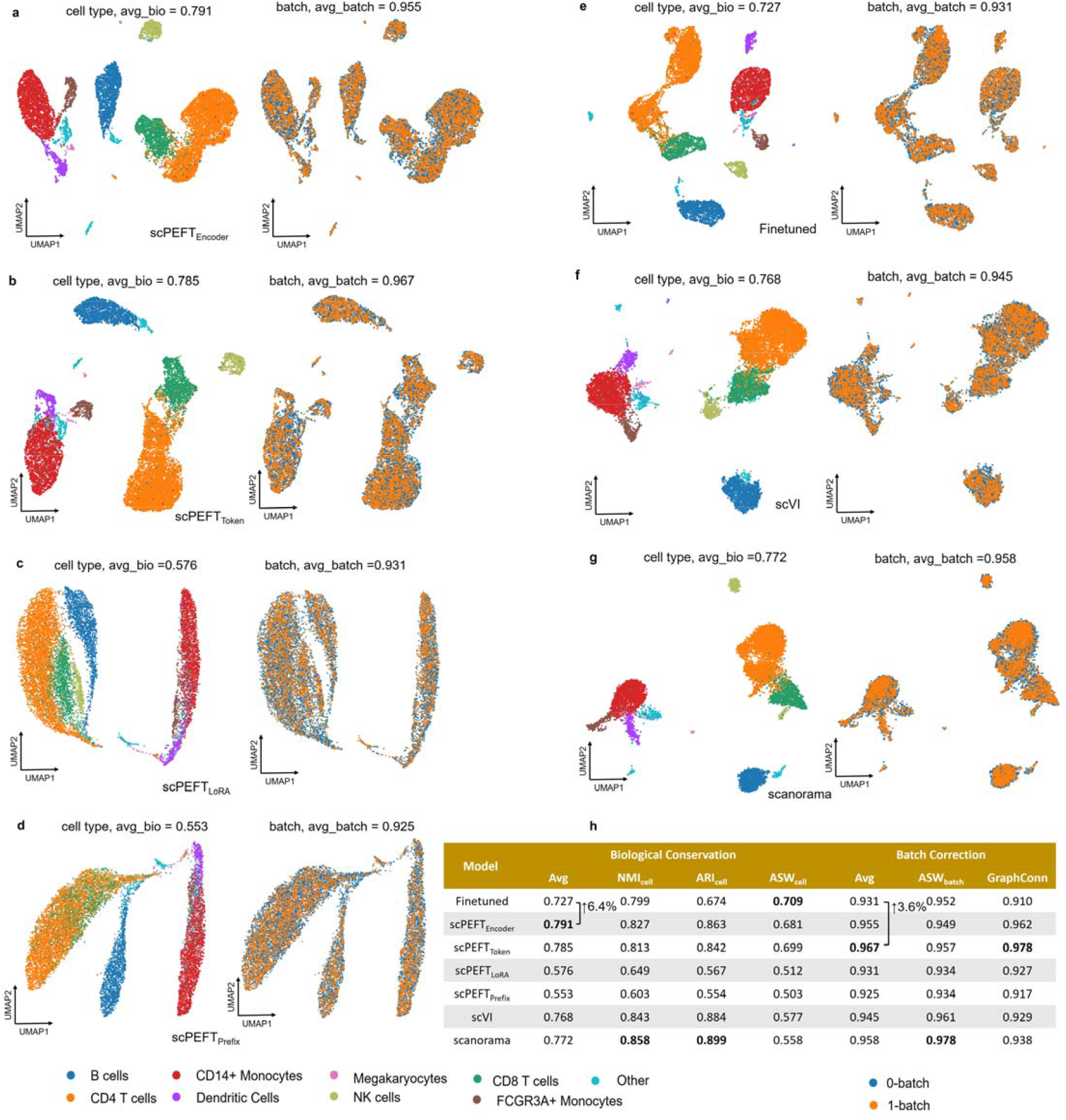
Batch correction results on the PBMC 10K dataset. **a-g**, UMAP visualizations of corrected cell embeddings, color-coded by cell type and batch, for scPEFT (using Encoder adapter, Token adapter, LoRA, and Prefix adapter), fine-tuned scGPT, scVI, and Scanorama. **h**, Comparative table of batch correction performance across these methods.

**Extended Data Figure 8:**
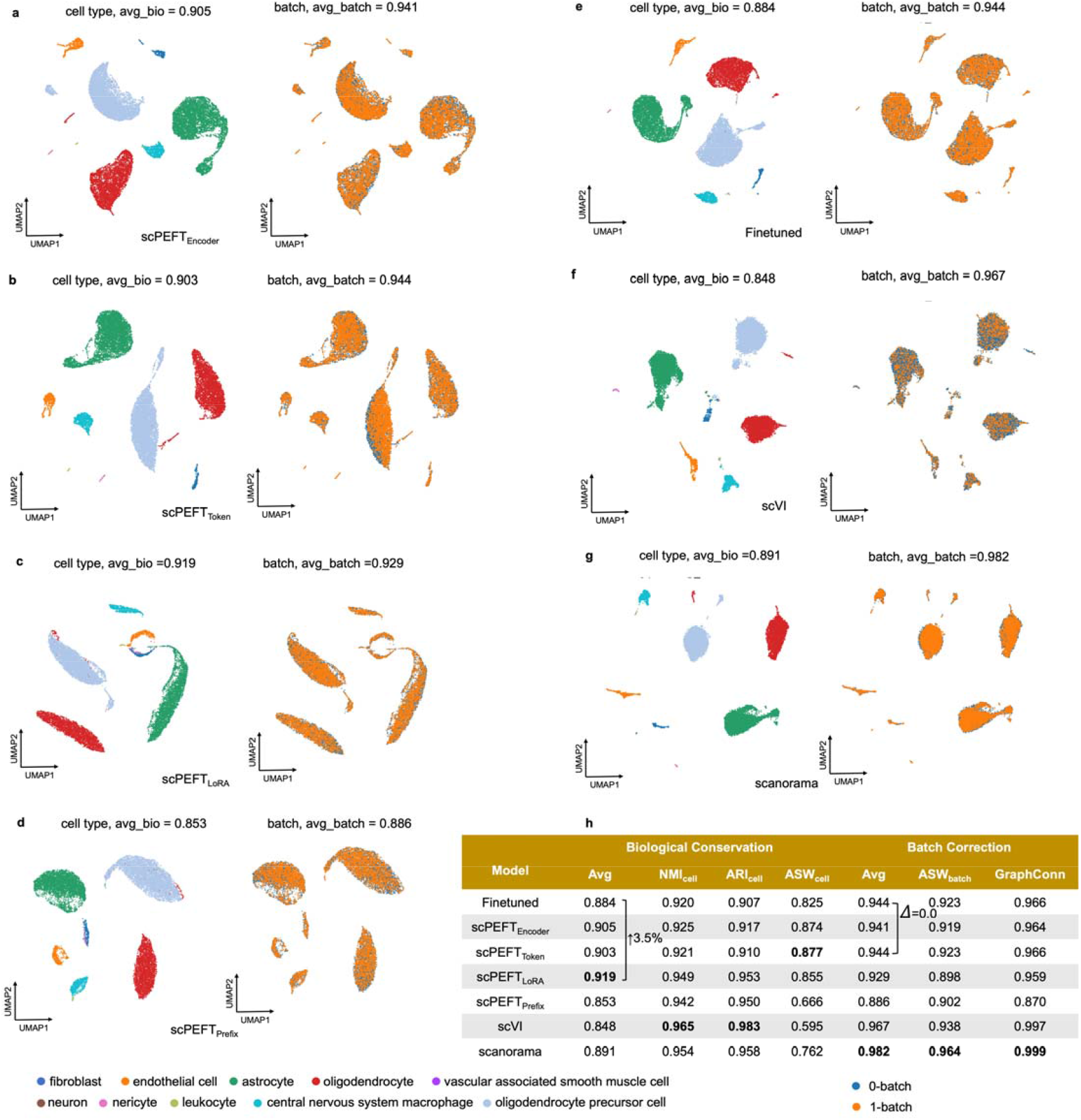
Batch correction results on the Perirhinal cortex dataset. **a-g**, UMAP visualizations of corrected cell embeddings, color-coded by cell type and batch, for scPEFT (using Encoder adapter, Token adapter, LoRA, and Prefix adapter), fine-tuned scGPT, scVI, and Scanorama. **h**, Comparative table of batch correction performance across these methods.

**Extended Data Figure 9:**
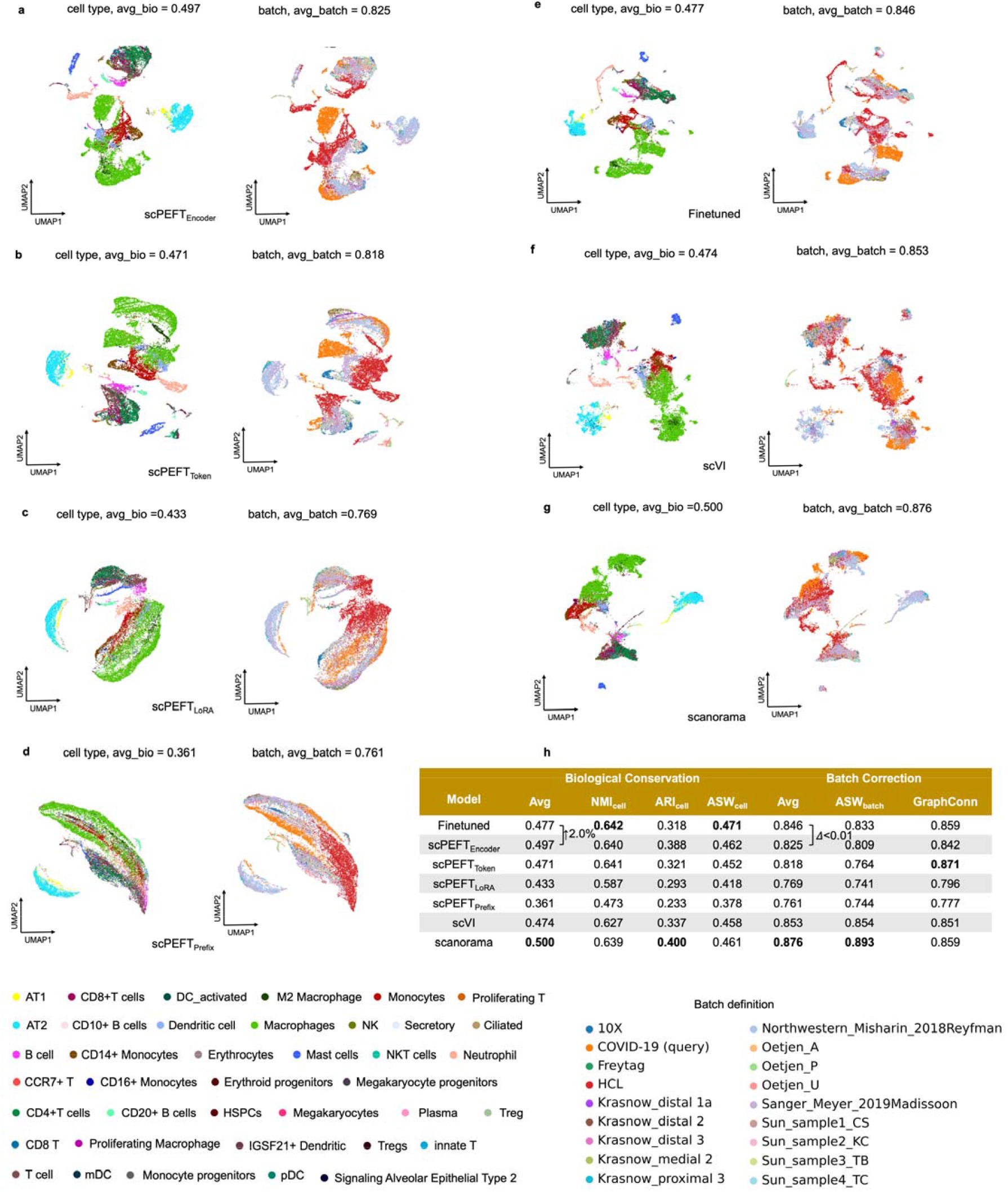
Batch correction results on the COVID-BATCH dataset. **a-g**, UMAP visualizations of corrected cell embeddings, color-coded by cell type and batch, for scPEFT (using Encoder adapter, Token adapter, LoRA, and Prefix adapter), fine-tuned scGPT, scVI, and Scanorama. **h**, Comparative table of batch correction performance across these methods.

**Extended Data Table 1:** Perturbation results on Adamson, Norman, Replogle_k562, and Replogle_rpe1 datasets.

|  | Adamson |  |  |  | Norman |  |  |  |
| --- | --- | --- | --- | --- | --- | --- | --- | --- |
|  | MSE | MSE_DE | PCC | PCC_DE | MSE | MSE_DE | PCC | PCC_DE |
| Finetuned scGPT | 0.00639 | **0.132** | 0.989 | **0.976** | 0.00421 | 0.226 | **0.987** | 0.863 |
| scPEFT<sub>Encoder</sub> | 0.00607 | 0.134 | **0.990** | 0.974 | **0.00420** | 0.277 | **0.987** | 0.851 |
| scPEFT<sub>Token</sub> | 0.00596 | 0.155 | **0.990** | 0.967 | 0.00495 | 0.318 | 0.984 | 0.829 |
| scPEFT<sub>LoRA</sub> | **0.00566** | 0.167 | **0.990** | 0.962 | 0.00498 | 0.322 | 0.984 | 0.829 |
| scPEFT<sub>Prefix</sub> | 0.00570 | 0.163 | **0.990** | 0.965 | 0.00497 | 0.321 | 0.984 | 0.829 |
| GEARS | 0.00729 | 0.262 | 0.988 | 0.952 | 0.00449 | **0.188** | **0.987** | **0.870** |

| Model | Replogle_k562 |  |  |  | Replogle_rpe1 |  |  |  |
| --- | --- | --- | --- | --- | --- | --- | --- | --- |
|  | MSE | MSE_DE | PCC | PCC_DE | MSE | MSE_DE | PCC | PCC_DE |
| Finetuned scGPT | **0.00699** | **0.134** | **0.987** | **0.935** | 0.0114 | **0.119** | **0.977** | 0.759 |
| scPEFT<sub>Encoder</sub> | 0.00727 | 0.143 | **0.987** | 0.933 | **0.0109** | 0.123 | **0.977** | 0.768 |
| scPEFT<sub>Token</sub> | 0.00720 | 0.151 | **0.987** | 0.931 | 0.0125 | 0.140 | 0.974 | **0.787** |
| scPEFT<sub>LoRA</sub> | 0.00747 | 0.163 | 0.986 | 0.930 | 0.0128 | 0.142 | 0.974 | 0.784 |
| scPEFT<sub>Prefix</sub> | 0.00706 | 0.153 | **0.987** | 0.931 | 0.0127 | 0.140 | 0.974 | 0.771 |
| GEARS | 0.00752 | **0.134** | 0.986 | 0.925 | 0.0138 | 0.132 | 0.971 | 0.705 |

