## Supplementary Figures and Tables for "Harnessing the Power of Single-Cell Large Language Models with Parameter Efficient Fine-Tuning using scPEFT"

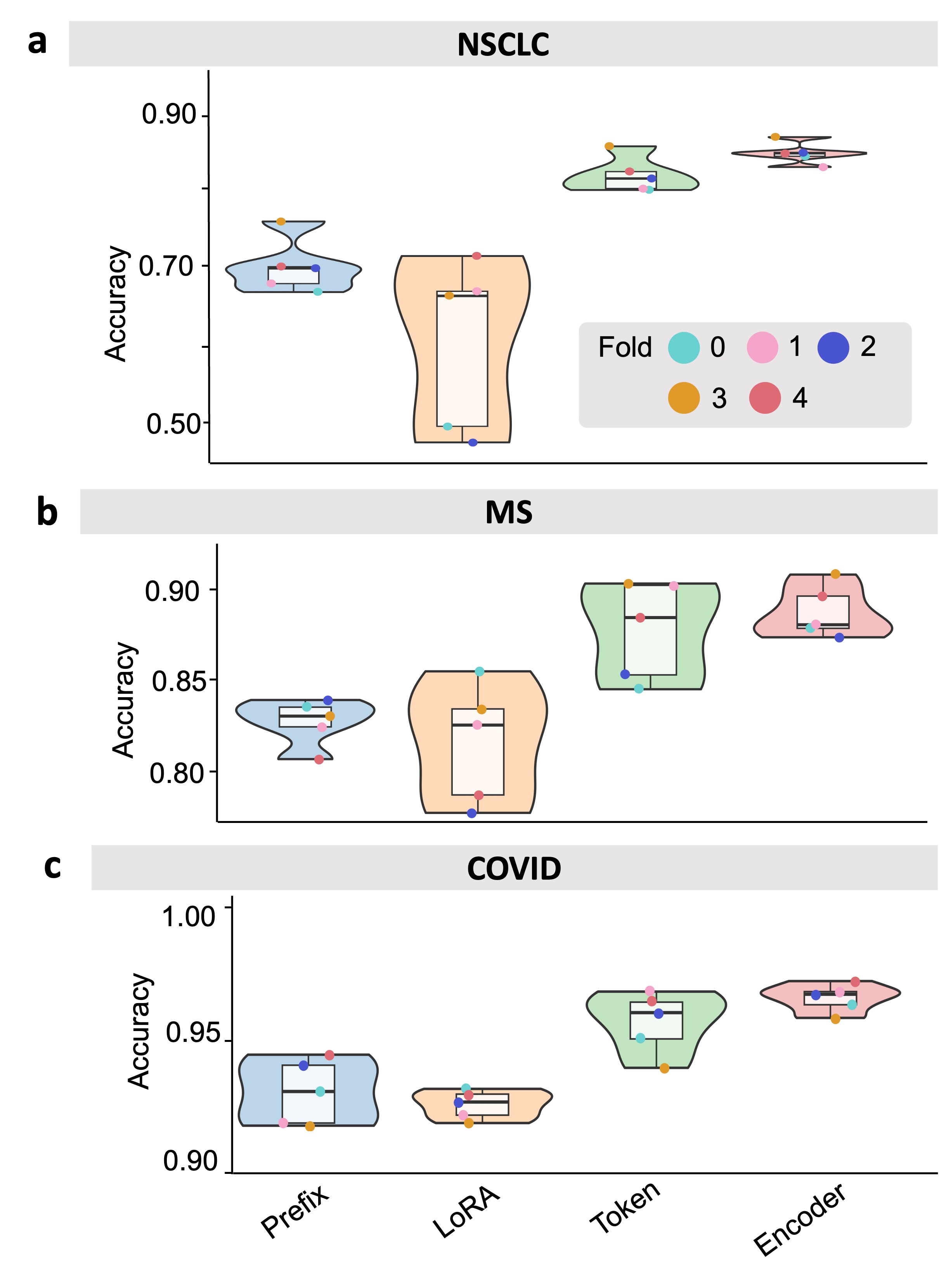

**Supplementary Figure 1**. Violin plots showcasing the performance of diverse adapters in scPEFT with the scGPT backbone for cell type identification across datasets: (**a**) NSCLC, (**b**) MS, and (**c**) COVID, respectively.

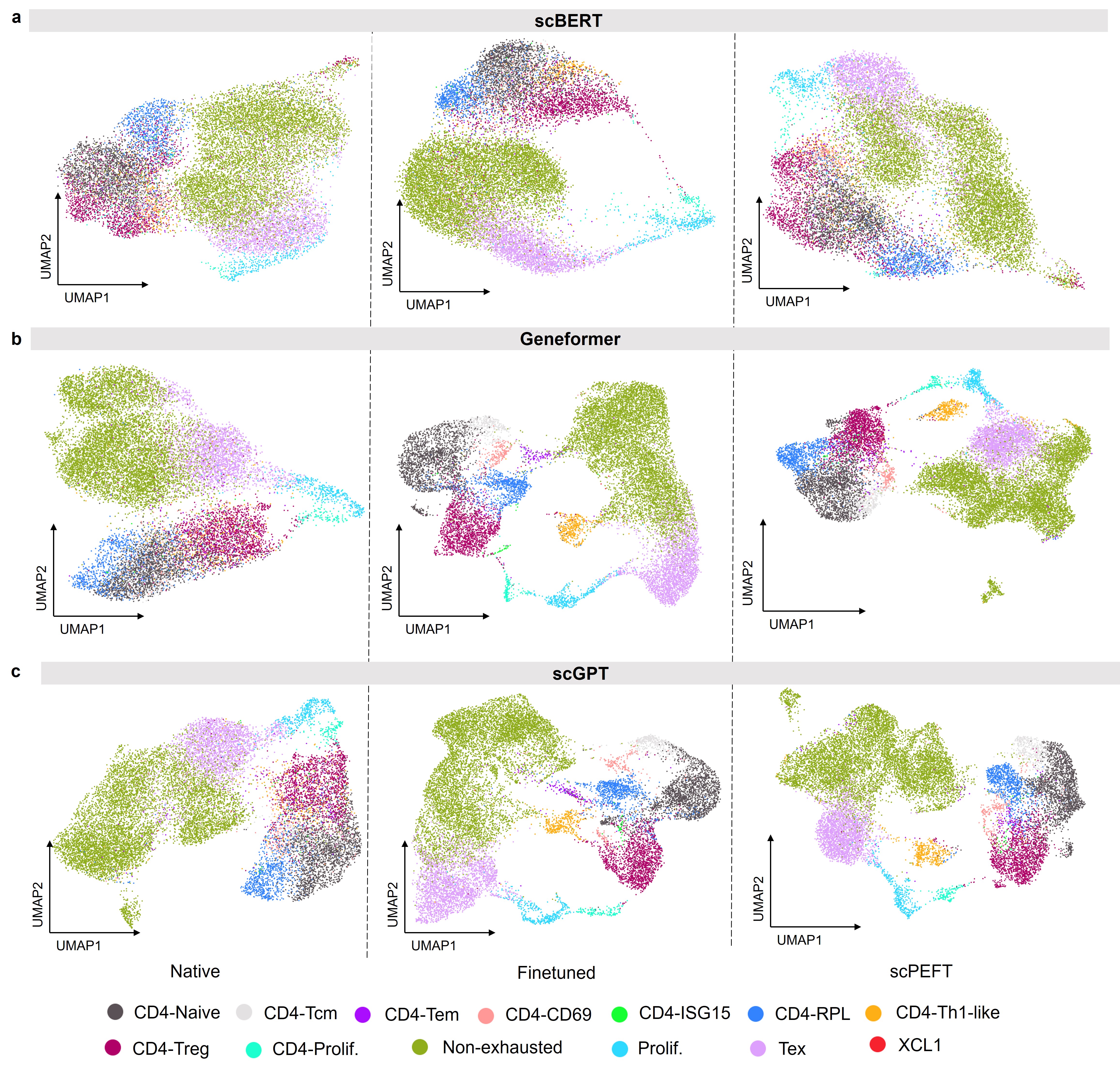

**Supplementary Figure 2.** UMAP visualizations of native, finetuned scLLMs, and scPEFT models with different backbones (**a**) scBERT, (**b**) Geneformer, and (**c**) scGPT, respectively, for cell type identification on the NSCLC dataset.

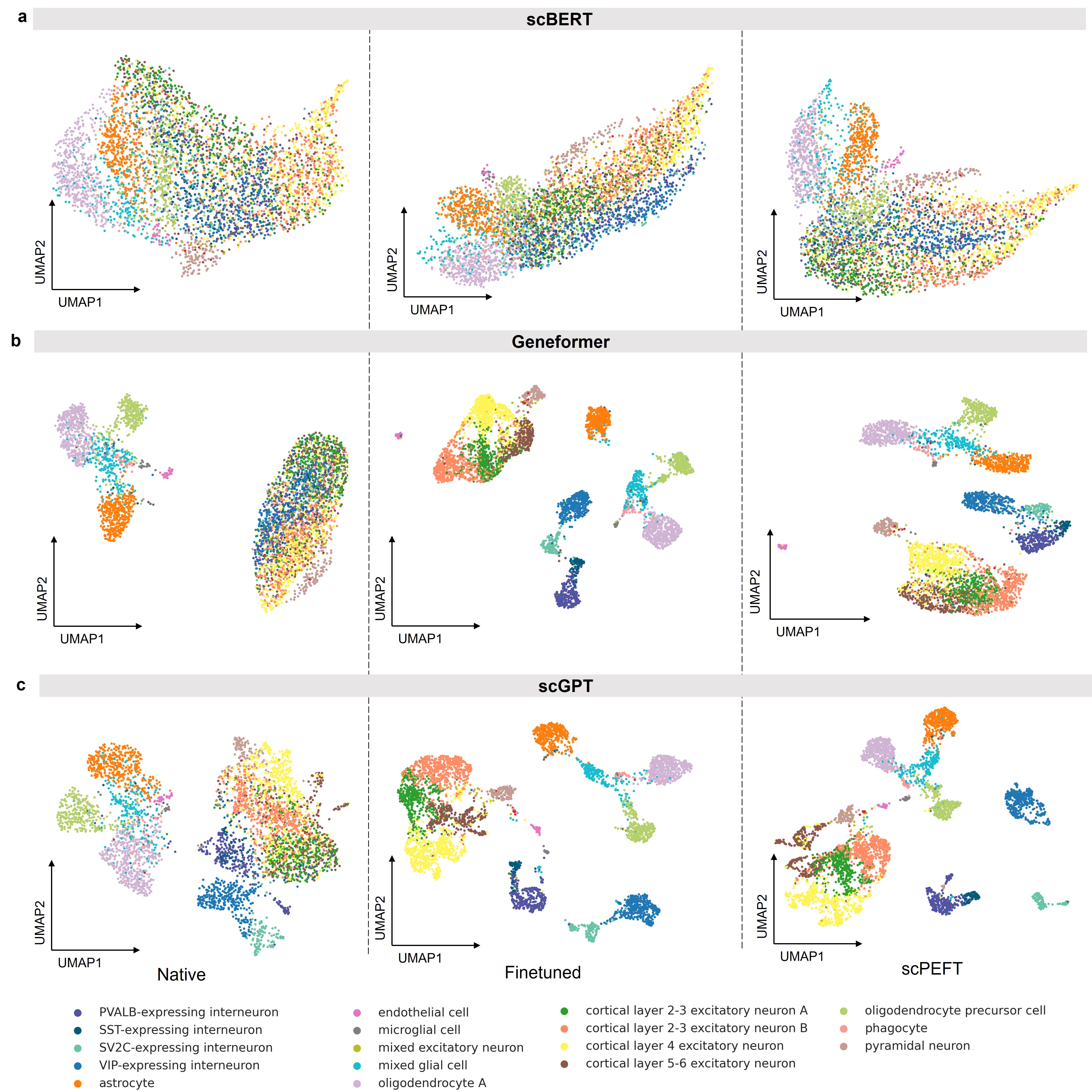

**Supplementary Figure 3.** UMAP visualizations of native, fine-tuned scLLMs, and scPEFT models with different backbones (**a**) scBERT, (**b**) Geneformer, and (**c**) scGPT, respectively, for cell type identification on the MS dataset.

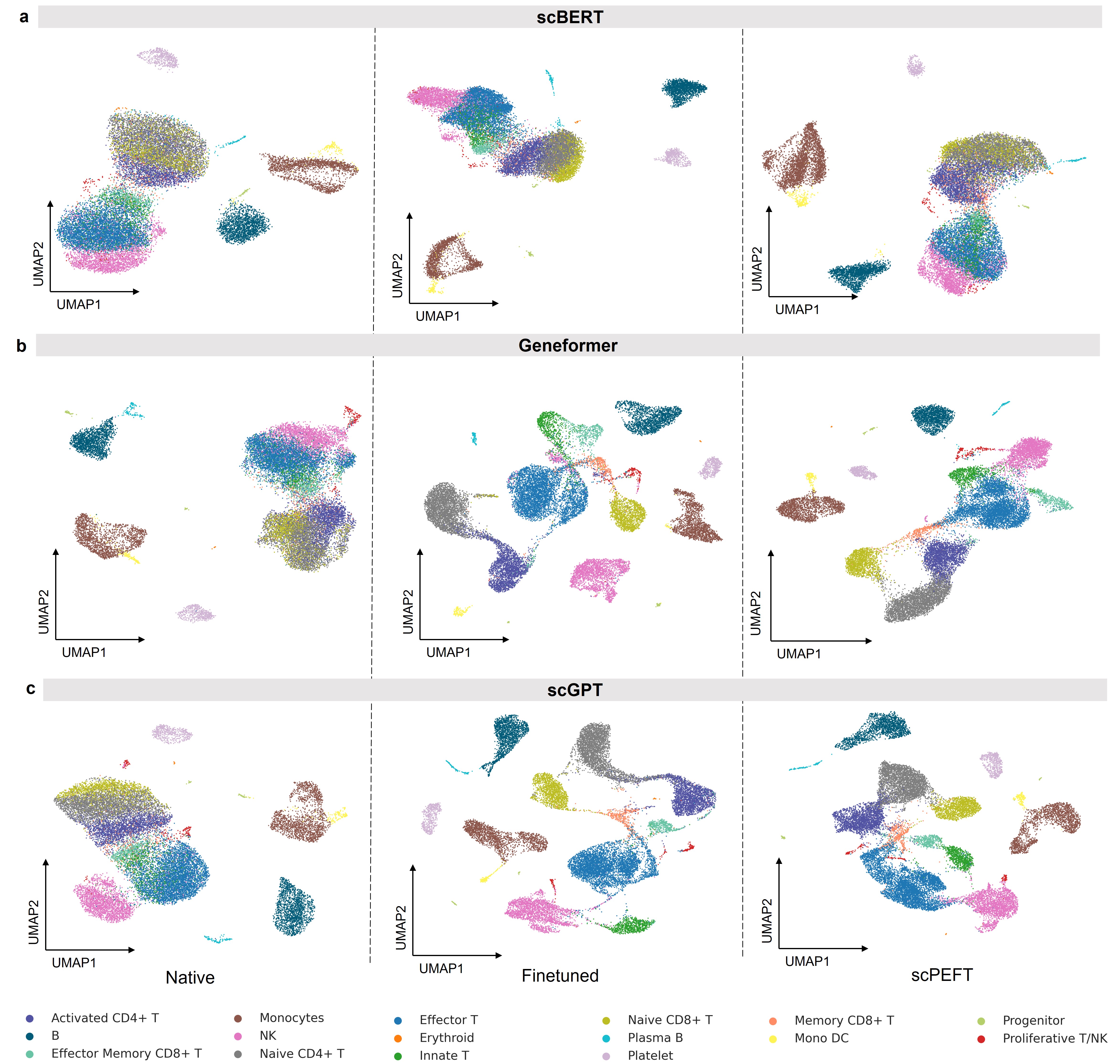

**Supplementary Figure 4.** UMAP visualizations of native, fine-tuned scLLMs, and scPEFT models with different backbones (**a**) scBERT, (**b**) Geneformer, and (**c**) scGPT, respectively, for cell type identification on the COVID dataset.

**Supplementary Table 1**. Default hyper-parameter settings

| **Hyperparameter** | | **Value** |
| --- | --- | --- |
| Token adapter | Fully connected layer 1 input dimension | 512 |
|  | Fully connected layer 1 output dimension | 128 |
|  | Fully connected layer 2 input dimension | 128 |
|  | Fully connected layer 2 output dimension | 512 |
|  | Activation function | GeLU |
|  | Residual connection | True |
| Encoder adapter | Fully connected layer 1 input dimension | 512 |
|  | Fully connected layer 1 output dimension | 128 |
|  | Fully connected layer 2 input dimension | 128 |
|  | Fully connected layer 2 output dimension | 512 |
|  | Activation function | GeLU |
|  | Residual connection | True |
|  | Adapter configuration to transformer layers | first 6 transformer blocks |
| Prefix adapter | Number of prefix tokens | 64 |
|  | Initial value of prefix token | 0 |
|  | Adapter configuration to transformer layers | first 6 transformer blocks |
| LoRA | LoRA rank | [**8**, 16, 32, 128, 256] |
|  | Lora Alpha | [**1**, 16, 32, 128, 256, 512] |
|  | LoRA input dimension | 512 |
|  | LoRA output dimension | 512 |
|  | Use LoRA on Q, K, and V, respectively | [True, False, True] |
|  | Adapter configuration to transformer layers | first 6 transformer blocks |
| Training | Optimizer | Adam |
|  | Optimizer momentum | β_1_ = 0.9, β_2_ = 0.999 |
|  | Maximum learning rate | 1e-5 |
|  | Ratio of epochs for learning rate schedule | 0.9 |
|  | Early stop patience | 5 |
|  | Maximum epoch | 50 |
|  | Batch size | 32 |
|  | Automatic mixed precision | True |
| scBERT  Geneformer  scGPT | Default adapter | Prefix adapter |
|  |  | Prefix adapter |
|  |  | Encoder adapter |
